# Emotional association enhances perceptual memory through amygdalo-cortical inputs during NREM sleep

**DOI:** 10.1101/2023.05.23.541852

**Authors:** Yoshihito Saito, Yuma Osako, Maya Odagawa, Yasuhiro Oisi, Chie Matsubara, Shigeki Kato, Kazuto Kobayashi, Mitsuhiro Morita, Joshua P. Johansen, Masanori Murayama

## Abstract

Emotional arousal is thought to enhance the consolidation of associated memories by activating the basolateral amygdala (BLA) and its projections to memory-storing regions^1–4^. Although the importance of both rapid eye movement (REM) and non-REM (NREM) sleep-state specific BLA activity for emotional memory processing has been proposed^5–9^, how and when the BLA interacts with other brain regions to enhance memory consolidation remains unclear^10^. Here, by adding emotional information to a perceptual recognition task that relies on top-down inputs from frontal to sensory cortices, we demonstrated that the BLA not only associates emotional information with perceptual information, but also enhances the retention of associated perceptual memory through BLA-frontal projections. Electrophysiological recordings revealed that emotional association increases the reactivation of coordinated activity across the BLA-frontal-sensory region during NREM sleep, but not during REM sleep. Notably, this inter-regional coordinated reactivation during NREM sleep was entrained to the BLA high-frequency oscillations in the emotional condition, suggesting that the BLA triggers inter-regional interaction. Optogenetic silencing of BLA terminals in the frontal cortex during NREM sleep, but not REM sleep, disrupted the enhanced retention of the perceptual memory, but not the association itself or the emotional component of associative memory. Our results indicate that the inter-regional coordination through the BLA-cortical inputs during NREM sleep is causally required for memory enhancement by emotional arousal.

---

Long-lasting memories of perceived information associated with emotionally arousing experiences (e.g., sexual, threatening) are critical for animal reproduction and survival^11^. Although human studies have shown that emotion-associated memories are retained longer than neutral memories^1, 12^, the underlying neural circuitry and physiological mechanisms remain unclear. Combining rodent behavioural tasks with neural circuit manipulation and high-spatiotemporal neural recordings has advantages for elucidating these mechanisms. However, because most rodent behavioural tasks inherently incorporate emotional information (e.g., water as a reward and foot shock as a threat), the behavioural outcomes of such tasks are already influenced by emotional information^13^. This makes it difficult to dissociate the effect of emotional information on memory processing of associated neutral information. To address this issue, we need to compare behavioural and neural data from behavioural tasks with and without emotional information.

The basolateral amygdala (BLA) is essential for associating neutral sensory information with emotional value, regardless of its valence^14, 15^, and has projections to many brain regions^16^. Thus, it has been proposed that the BLA modulates memory consolidation processes that take place in other brain regions^2–4^. However, how the BLA interacts with other brain regions via its direct projections to enhance memory consolidation remains unclear. We have previously reported that neutral perceptual (somatosensory) memory in mice is consolidated by top-down inputs from the secondary motor cortex (M2) to the primary somatosensory cortex (S1), independent of the hippocampus^17^. Therefore, we hypothesised that if perceptual memory is enhanced by emotional association, the enhancement would occur via BLA projections to one or both of these cortical regions.

Memory consolidation is closely linked to sleep state-dependent neural activity^18, 19^. While previous studies have suggested that rapid eye movement (REM) sleep-specific BLA activity is associated with emotional memory processing^5, 6, 13^, recent studies have highlighted the importance of non-REM (NREM) sleep-specific activity^7–9^. Despite the large body of research in this area, causal evidence is still lacking to resolve the controversy over a sleep-state-specific role of the BLA in memory modulation.

Here, using both a non-emotional perceptual recognition task and a perceptual-emotional associative recognition task, we show that emotional information enhances the retention of associated perceptual information. Anatomical and chemogenetic functional circuit analyses reveal that M2- projecting BLA neurons are required for both perceptual-emotional association and enhanced retention of perceptual memory. Multi-site electrophysiological recordings and sleep-state-dependent optogenetic silencing demonstrate that BLA-M2 inputs during NREM, but not REM, sleep are necessary for enhanced retention of perceptual memory.

## Emotional association enhances perceptual memory

To assess the effect of the emotional association on the associated emotionally neutral perceptual memory, we employed three behavioural tasks. First, we used the perceptual recognition task developed in our previous study^17^ to assess the recognition memory for floor texture as an emotionally neutral control (Fig. 1a). In this task, male mice underwent learning sessions in an arena containing two empty cups with smooth floors, followed by a 24-hour rest period in their home cage. On day 2, the mice were presented with an arena in where one floor had been replaced with a novel grooved texture and were given 10 minutes to explore as a recall session. Consistent with our previous finding^17^ and innate rodent behaviour, the mice showed a preference for the cup on the novel texture (Fig. 1d left, without female). Second, we developed an “associative recognition task” to assess the effect of emotional information on the texture memory by providing a female mouse as a natural reward for male mice (Fig. 1b and Extended Data Fig. 1)^20, 21^. In the learning sessions of this task, male mice were presented with an area containing a female mouse in one of the two cups on smooth floors. After the resting period, memory recall sessions were performed as with the perceptual recognition task on day 2. In contrast to the perceptual recognition task, the mice that had experienced both the texture and the female showed a preference for the cup on the familiar texture during the memory recall sessions (Fig. 1d left, with female) and an increased total contact time with both cups (Fig. 1d right). This reversal of the texture preference and increased total contact time may reflect changes in exploration strategy resulting from the association between the texture and the female mouse. To determine whether the discrimination between the novel and familiar floor texture is based on the recognition of texture information, and whether the increased total contact time is independent of texture information, we conducted the recall sessions of the associative recognition task in an arena with familiar smooth floors on both sides (Fig. 1c, with female: same texture). In this condition, the mice did not show a preference during the recall sessions (Fig. 1d left, with female: same texture), indicating that the mice used texture information to discriminate the female-presented side rather than other sensory cues (e.g., visual, olfactory). However, they still showed an increased total contact time (Fig. 1d right). These results indicate that while the texture-female association reverses the sign of texture preference during recall sessions, discrimination between the novel and familiar textures reflects the texture memory, and increased total contact time reflects the emotional component (i.e., female memory) of associative memory.

**Fig. 1.**
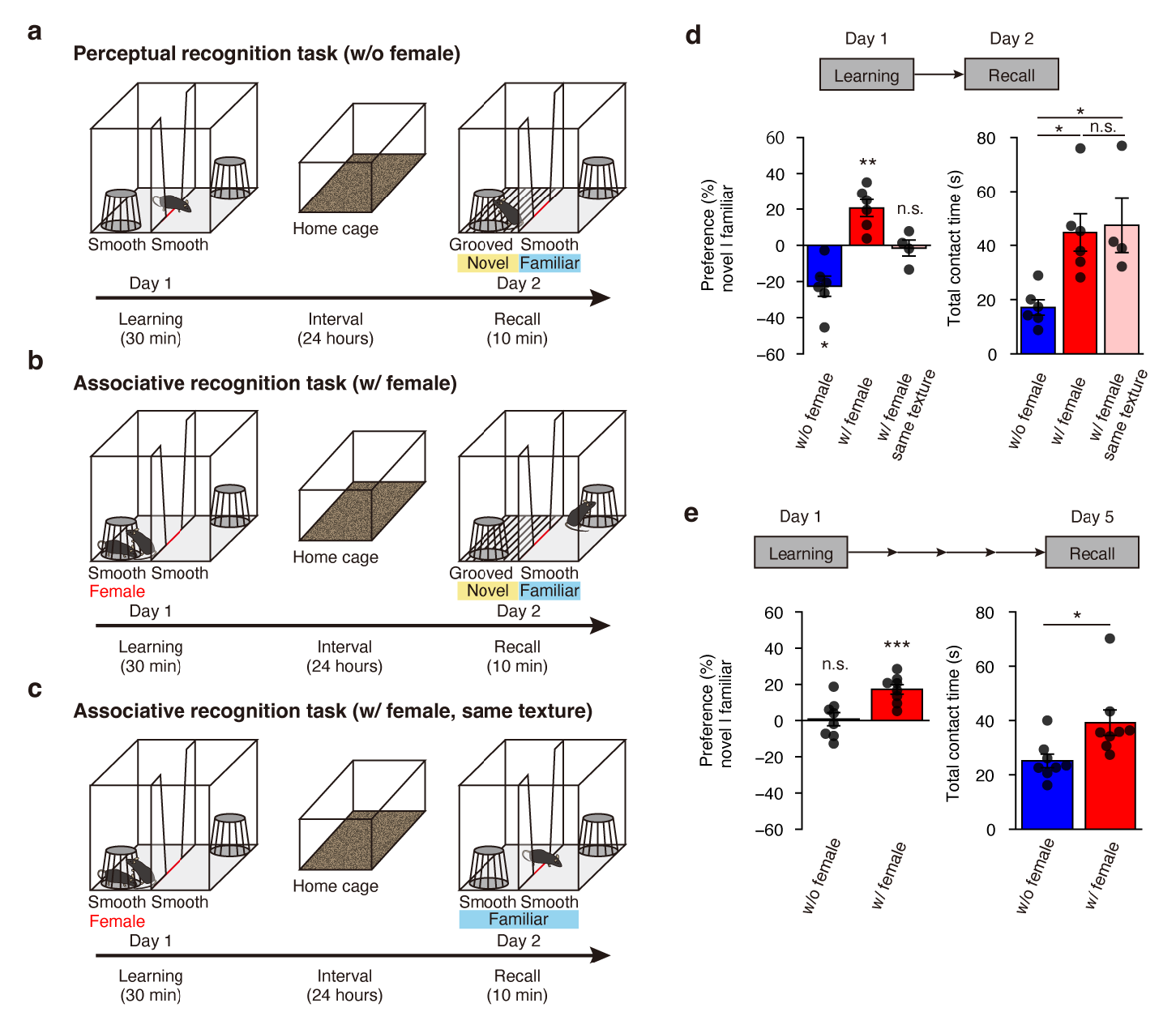
Female presentation enhances texture memory retention in male mice. **a**, Schematics of perceptual recognition task. **b**, Schematics of associative recognition task. **c**, Schematics of same texture control of the associative recognition task. **d**, Memory performance during the recall sessions on day 2 (w/o female, *n* = 6; w/ female, *n* = 6; w/ female (same texture), *n* = 4). Preferences of exploratory behaviour (w/o female, *t*_(5)_ = −4.01, * *P* = 0.0101; w/ female, *t*_(5)_= −4.35, ** *P* = 0.0074; w/ female (same texture), *t*_(3)_ = −0.35, n.s. *P* = 0.7466, one-sample t-test against chance level) (left). Total contact time with the cups on both sides (F(2, 13) = 7.02, * *P* < 0.05, n.s. *P* > 0.05, one-way analysis of variance with Tukey’s post-hoc test) (right). **e**, Memory performance during the recall sessions on day 5 (w/o female, *n* = 8; w/ female, *n* = 8). Preferences of exploratory behaviour (w/o female, *t*_(7)_ = 0.22, n.s. *P* = 0.8353; w/ female, *t*_(7)_ = 6.44, *** *P* = 0.0004, one-sample t-test against chance level) (left). Total contact time with the cups on both sides (*t*_(14)_ = −2.63, * *P* = 0.0198, Student’s t-test) (right). Data are presented as mean ± SEM.

To test whether association with the female enhances the retention of texture memory, we extended the resting period from 24 to 96 hours and conducted recall sessions for both the perceptual and associative recognition tasks on day 5. The results showed that the mice in the perceptual recognition task showed no texture preference in this condition, whereas mice in the associative recognition task retained their preference for the familiar texture and increased their total contact time (Fig. 1e). These results indicate emotional association enhances the retention of the associated perceptual memory.

## Anatomical and molecular features of the BLA-cortical circuit

We next explored which neural circuits underlie the enhanced retention of the texture memory by female association. We first examined the BLA activation in male mice by the female presentation using immunohistological analysis of c-Fos expression (Extended Data Fig. 2a). Consistent with a previous report^22^, we confirmed that the female presentation increased the density of c-Fos+ cells in the BLA (Extended Data Fig. 2b,c). Since our previous study showed that top-down inputs from the secondary motor cortex (M2) to the primary somatosensory cortex (S1) are essential for texture memory consolidation^17^, we investigated the connections among the BLA, M2 and S1. Labelling of axonal projections from the BLA by injecting an adeno-associated virus (AAV) vector expressing green fluorescent protein (AAV1-hSyn-GFP) into the BLA as an anterograde tracer (Fig. 2a,b) revealed strong BLA projections to frontal cortical regions including the M2 (Fig. 2c,d) and weak ones to the S1 (Fig. 2e). Based on these results, we examined whether the M2 is the hub region connecting the BLA and S1. To visualise neurons that receive inputs from the BLA and send outputs to the S1, we injected AAV1-hSyn-Cre into the BLA, which infects in an anterograde-transsynaptic manner and expresses Cre recombinase in post-synaptic neurons^23^. We also injected AAVrg-CAG-FLEX-tdTomato into the S1 to selectively label S1-projecting neurons expressing Cre recombinase with a red fluorescence protein (Fig. 2f,g). Although BLA axons were distributed widely in frontal cortices (Fig. 2c), BLA-recipient S1-projecting neurons were predominantly in the M2 (Fig. 2h-j).

**Fig. 2.**
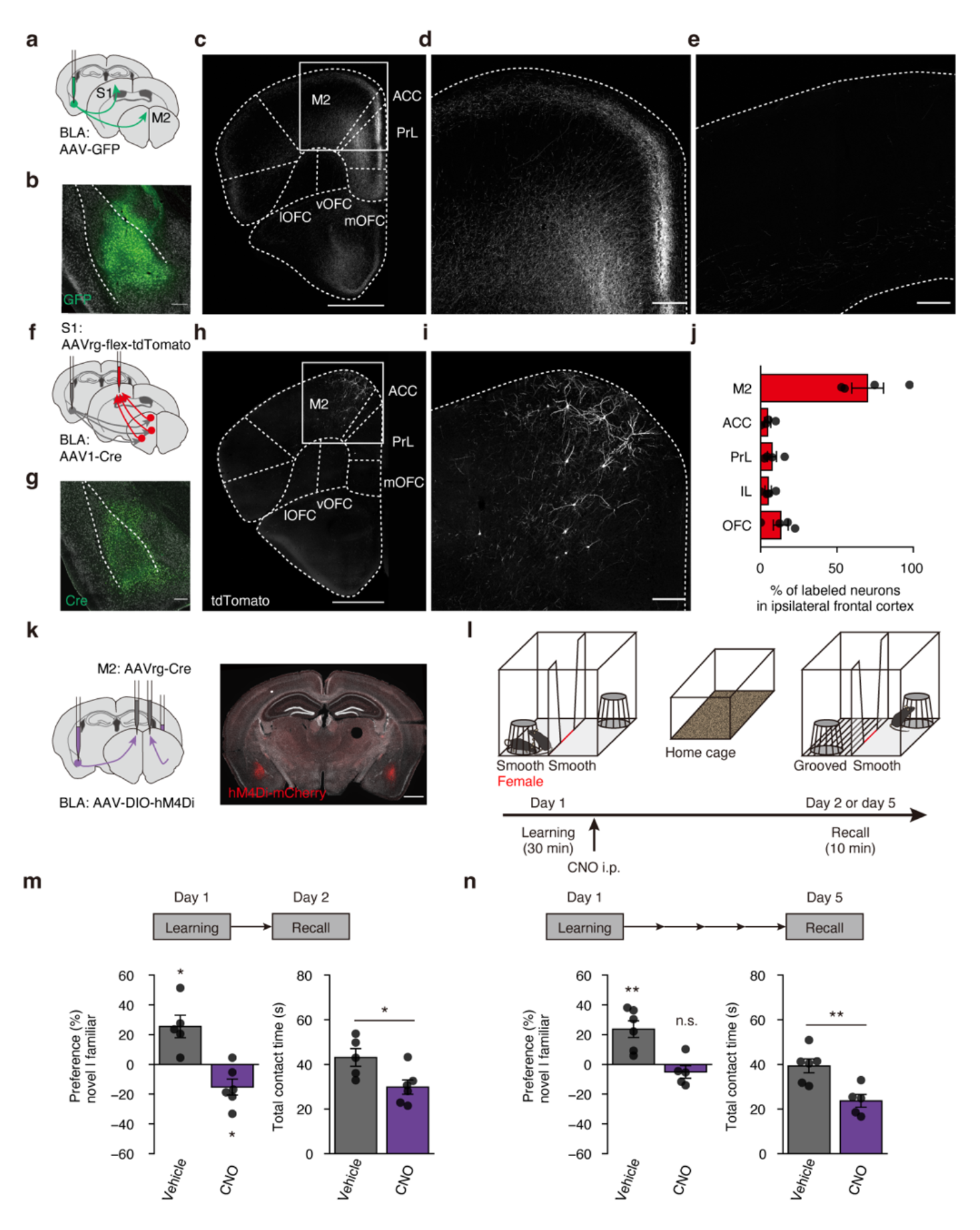
M2-projecting BLA neurons not only store the female memory, but also enhance the texture memory consolidation. **a**, Anterograde tracing of BLA axons. **b**, AAV-GFP was unilaterally injected into the BLA (scale bar, 200 μm). **c**, Labelled BLA axons were distributed in the frontal cortical regions (scale bar, 1 mm). Secondary motor cortex, M2; Anterior cingulate cortex, ACC; Prelimbic cortex, PrL; infralimbic cortex, IL; lateral orbitofrontal cortex, lOFC; ventral OFC, vOFC; medial OFC, mOFC. **d**, Magnified view of the area indicated in (c) showing dense innervation in M2 by BLA axons (scale bar, 200 μm). **e**, BLA axons were not observed in the S1 (scale bar, 200 μm). **f**, Labelling BLA-recipient S1-projecting neurons. AAV1-hSyn-Cre and AAVrg-flex-tdTomato were unilaterally injected into the BLA and S1, respectively. **g**, BLA injection site (scale bar, 200 μm) confirmed by expression of Cre. **h**, Labelled neurons in frontal cortical regions (scale bar, 1 mm). **i**, Magnified view of the area indicated in (**h**) (scale bar, 200 μm). **j**, Distribution of BLA-recipient S1- projecting neurons in the ipsilateral frontal cortex (*n* = 4). **k**, Inactivation of the M2-projecting BLA neurons by chemogenetics. AAVrg-Cre and AAV-DIO-hM4Di were bilaterally injected into the M2 and BLA, respectively (left). Expression of hM4Di was localized in the BLA (scale bar, 1 mm) (right). **l**, CNO or vehicle was injected immediately after learning sessions of the associative memory task, and then recall sessions were performed on day 2 or day 5. **m**, Memory performance during the recall sessions on day 2 (vehicle, *n* = 5; CNO, *n* = 6). Preferences of exploratory behaviour (vehicle, *t*_(4)_ = 3.37, * *P* = 0.0280; CNO, *t*_(5)_ = −2.84, * *P* = 0.0364, one-sample t-test against chance level) (left). Total contact time with the cups on both sides (*t*_(9)_ = 2.64, * *P* = 0.0270, Student’s t-test) (right). **n**, Memory performance during the recall sessions on day 5 (vehicle, *n* = 6; CNO, *n* = 5). Preferences of exploratory behaviour (vehicle, *t*_(5)_ = 4.22, ** *P* = 0.0083 CNO, *t*_(4)_ = −1.20, n.s. *P* = 0.2959, one-sample t-test against chance level) (left). Total contact time with the cups on both sides (*t*_(9)_ = 3.67, ** *P* = 0.0051, Student’s t-test) (right). Data are presented as mean ± SEM.

A recent study showed that BLA neurons expressing *Fezf2* encode either positive or negative valence and are necessary for reward or punishment-based learning via projections to the olfactory tubercle (OT) and nucleus accumbens (NAc), respectively^24^. Based on this knowledge, we investigated whether M2-projecting BLA neurons exhibit these properties. Fluorescence *in situ* hybridisation for *Fezf2* in mice injected with AAVrg-CAG-H2B-GFP into the M2 showed that most M2-projecting BLA neurons (79.64±1.5%) expressed *Fezf2* (Extended Data Fig. 3a-d). Furthermore, we observed axonal projections of M2-projecting BLA neurons by injecting AAVrg-pgk-Cre and AAV1-EF1a-DIO-EYFP into the M2 and BLA, respectively, and found that the axon collaterals of M2-projecting BLA neurons were widely distributed in frontal cortical and striatal regions, including the OT and NAc (Extended Data Fig. 3e-f). Together, these results suggest that M2-projecting BLA neurons send emotional valence information to various brain regions and may influence the M2-S1 top-down circuit underlying texture memory consolidation.

**Fig. 3.**
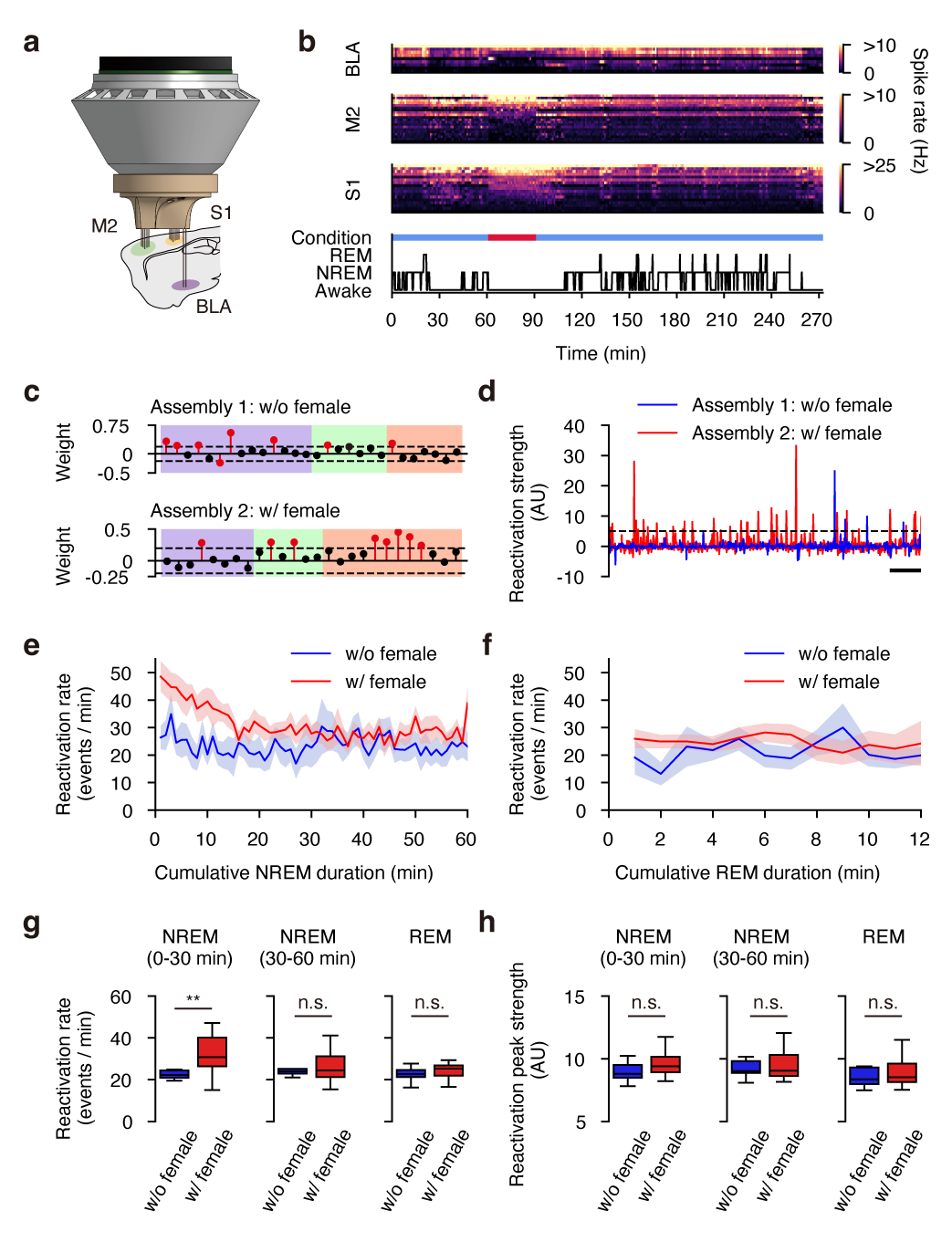
Female presentation increases the reactivation rate of tri-regional assemblies during post-NREM, but not post-REM. **a**, Simultaneous extracellular recording from the BLA, M2, and S1. **b**, Activity of simultaneously recorded units and hypnogram throughout pre-learning epoch (60 min), learning epoch (30 min), and post-learning epoch (180 min). Spike rates were calculated in a 1 min time bin. **c**, Example weight vectors of tri-regional assemblies in the without (top) and with (bottom) female group. Tri-regional assemblies were defined by whether the assembly contained at least single member units (red lollipops) from each region that exceeded the threshold (dashed line). **d**, Examples of traced reactivations of the assemblies (**c**). Reactivation events were detected when the strength exceeded the threshold 5 (dashed line). **e**, Time-aligned mean reactivation rates of tri-regional assemblies during post-NREM (red, with female, *n* = 17 assemblies; blue, without female, *n* = 9 assemblies; shade represents 95% confidence interval). **f**, Same layout as in (**e**) for post-REM. **g**, Reactivation rates during the first 30 min of post-NREM (left, 0-30 min), second 30 min of post-NREM (middle, 30-60 min) and post-REM (right) (NREM 0-30 min, ** *P* = 0.0021; NREM 30-60 min, n.s. *P* = 0.2766; REM, n.s. *P* = 0.1288, Mann-Whitney U test). **h**, Peak strength of reactivation events during first 30 min of post-NREM (left, 0-30 min), second 30 min of post-NREM (middle, 30- 60 min) and post-REM (right) (NREM 0-30 min, n.s. *P* = 0.0728; NREM 30-60 min, n.s. *P* = 0.4571; REM, n.s. *P* = 0.1660, Mann-Whitney U test). Boxplots show the median with 25/75 percentile (box) and 1.5 × interquartile range (whiskers).

## Memory enhancement requires BLA-cortical circuit

To examine the functional role of M2-projecting BLA neurons in the texture-female associative memory, we silenced these neurons by chemogenetics. AAVrg-pgk-Cre and AAV8-hSyn-DIO-hM4Di were injected into the bilateral M2 and BLA, respectively, to express an inhibitory Gi-DREADD (designer receptors exclusively activated by designer drugs) in M2-projecting BLA neurons (Fig. 2k). After learning sessions of the associative recognition task, clozapine N-oxide (CNO) was injected intraperitoneally to silence hM4Di expressing neurons (Fig. 2l). On day 2 recall sessions, CNO-treated mice showed a preference for the novel texture and a decrease in total contact time compared with the control mice (Fig. 2m). Since it was assumed that if this chemogenetic silencing only suppressed the consolidation of the female memory component of associative memory, mice would seek the novel texture side based on innate preference and the total contact time would decrease, these results indicate that M2-projecting BLA neurons are necessary for the consolidation of the female memory, but not for the baseline texture memory consolidation. Furthermore, we examined the contribution of M2- projecting BLA neurons to the enhanced retention of the texture memory by conducting memory recall sessions on day 5. Although the control mice showed a preference for the familiar side, CNO-treated mice showed no texture recognition and decreased total contact time (Fig. 2n), indicating that M2- projecting BLA neurons not only store the female memory component of associative memory, but also enhance the texture memory consolidation.

## Emotional experience gains BLA-cortical reactivation

To begin to understand when and how the BLA interacts with the cortical circuit to enhance texture memory consolidation during sleep, we first asked whether the female presentation changes the proportion of time spent in the awake, NREM, and REM states by classifying those with electroencephalogram (EEG) and electromyogram (EMG) recordings (Extended Data Fig. 4a,b). We found no apparent alterations in the sleep architecture upon the female presentation (Extended Data Fig. 4c). Next, to explore how neural activity of the BLA, M2, and S1 during sleep after learning epochs (post-NREM or post-REM) contributes to the texture memory enhancement, we recorded single unit spiking from these regions and examined the sleep-dependent spike rate changes in either with or without female conditions (Fig. 3a,b and Extended Data Fig. 5a, and Supplementary Table 1). In contrast to M2 and S1 units, BLA units increased spike rate during both post-NREM and post-REM, compared with learning epochs in both conditions (Extended Data Fig. 5b)^8^. However, we observed no changes in the spike rate ratio upon the female presentation in these areas (Extended Data Fig. 5c).

**Fig. 4.**
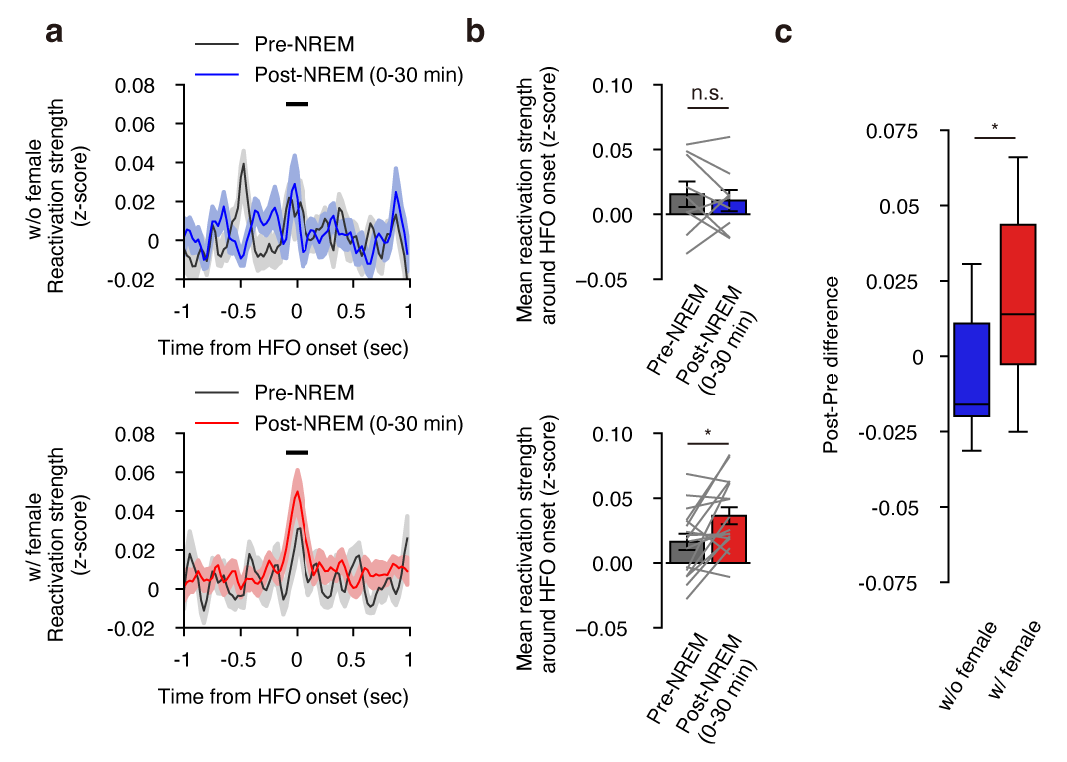
Female presentation enhances the inter-regional reactivation around BLA-HFOs. **a**, Mean z-scored reactivation strength of tri-regional assemblies around BLA-HFOs onset in without female condition (top) and with female condition (bottom) during pre-NREM and first 30 min of post-NREM (shade represents ± SEM). **b**, Mean reactivation strength in a ± 100-ms window around BLA-HFOs onset in without female condition (top) and with female condition (bottom) during pre-NREM and post-NREM (0-30 min) (without female, n.s. *P* = 0.4961; with female, * *P* = 0.0150, Wilcoxon signed rank-test). **c**, The pre-NREM and post-NREM (0-30 min) difference in mean reactivation strength around the onset of BLA-HFOs was compared in the without female (blue) and with female (red) conditions (* *P* = 0.0155, Mann-Whitney U test). Boxplots show the median with 25/75 percentile (box) and 1.5 × interquartile range (whiskers).

**Fig. 5.**
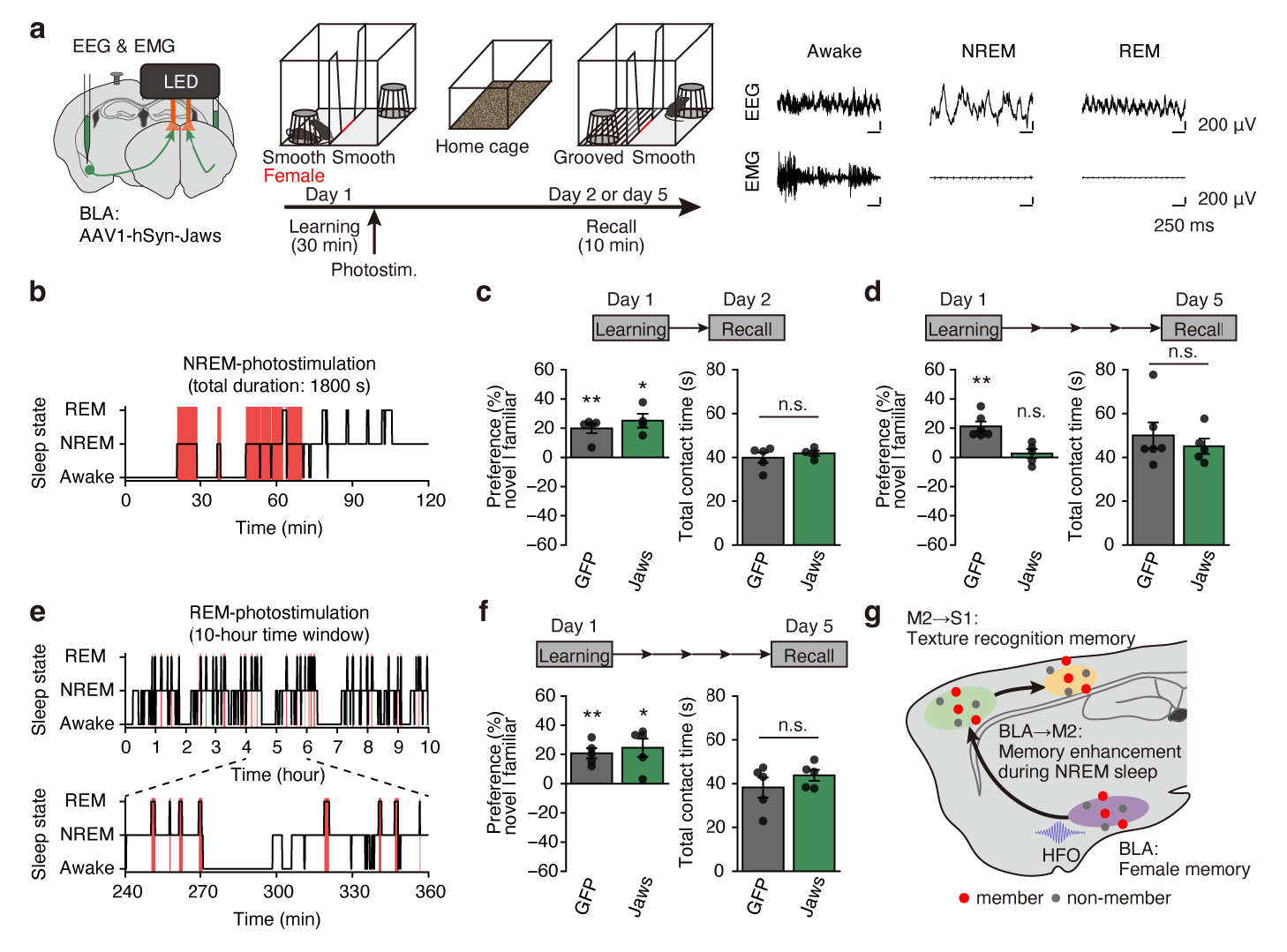
BLA-M2 inputs during post-NREM, but not post-REM, are required for the enhancement of the texture memory consolidation. **a**, Sleep-state dependent silencing of BLA axons at M2 by optogenetics. AAV-Jaws was bilaterally injected into the BLA and optic fibbers were placed on the M2. Photostimulation was delivered during post-NREM (total duration: 30 min) or post-REM sleep (10-h time window) of learning sessions of the associative memory task, then recall sessions were performed on day 2 or 5 (left). Sleep states were classified online by EEG and EMG (right). **b**, An example of sleep states and post-NREM sleep specific photostimulation (shaded red bar). **c**, Memory performance during the recall sessions on day 2 in opto-NREM condition (GFP, *n* = 5; Jaws, *n* = 4). Preferences of exploratory behaviour (GFP, *t*_(4)_ = 6.02, ** *P* = 0.0038; Jaws, *t*_(3)_ = 5.33, * *P* = 0.0129, one-sample t-test against chance level) (left). Total contact time with the cups on both sides (*t*_(7)_ = −0.72, n.s. *P* = 0.4974, Student’s t-test) (right). **d**, Memory performance during the recall sessions on day 5 in opto-NREM condition (GFP, *n* = 6; Jaws, *n* = 5). Preferences of exploratory behaviour (GFP, *t*_(5)_ = 6.54, ** *P* = 0.0012; Jaws, *t*_(4)_ = 0.84, n.s. *P* = 0.4465, one-sample t-test against chance level) (left). Total contact time with the cups on both sides (*t*_(9)_ = 0.67, n.s. *P* = 0.5180, Student’s t-test) (right). **e**, An example of sleep states and post-REM sleep specific photostimulation (shaded red bar). **f**, Memory performance during the recall sessions on day 5 in opto-REM condition (GFP, *n* = 5; Jaws, *n* = 5). Preferences of exploratory behaviour (GFP, *t*_(4)_ = 6.04, ** *P* = 0.0038; Jaws, *t*_(4)_ = 3.95, * *P* = 0.0168, one-sample t-test) (left). Total contact time with the cups on both sides (*t*_(8)_ = −1.07 n.s. *P* = 0.3161, Student’s t-test) (right). **g**, Schematic summary of proposed mechanism of the enhancement of the texture memory consolidation by female presentation. Texture memory and female memory are consolidated in the M2-S1 circuit and in the BLA, respectively, and increased inter-regional reactivations through BLA-M2 inputs during post-NREM contribute to the enhancement of the texture memory consolidation. Data are presented as mean ± SEM.

A growing body of evidence suggests that sleep-dependent reactivations of learning-related coordinated activity patterns (neural assemblies) lead to memory consolidation^8, 9, 17, 25, 26^. Therefore, we investigated whether female presentation enhances reactivations of neural assemblies across the BLA, M2, and S1 (tri-regional assembly) during sleep. We applied a principal component analysis and independent component analysis (PCA/ICA) to neural activity during the learning epoch to identify tri-regional assemblies and calculated the reactivation strength of these assemblies during post-NREM and post-REM (Fig. 3c,d and Extended Data Fig. 6, and Supplementary Table 1)^27, 28^. We ensured the inter-regional interactions during reactivations by calculating the reactivation window specific cross-correlogram of BLA-M2, M2-S1, and BLA-S1 unit pairs in the same assemblies (member unit) (Extended Data Fig. 7). Notably, female presentation increased reactivation rates of the tri-regional assemblies during post-NREM immediately after learning, and the reactivation rates gradually decreased over the first hour of post-NREM (Fig. 3e,g). In contrast, we observed no significant difference in either the reactivation rates during post-REM sleep (Fig. 3f,g) or in the peak strengths of the reactivation events during post-NREM and post-REM upon female presentation (Fig. 3h). These results indicate that female presentation increases reactivations of learning-related activity across BLA, M2 and S1 specifically during early post-NREM sleep.

To further understand the role of the BLA during NREM sleep, we focused on high-frequency oscillations (HFOs) observed in the BLA^9, 29, 30^. HFOs are generated locally in the BLA^30^ and are associated with the synchronous spiking activity of BLA neurons (Extended Data Fig. 8a-d). Thus, these HFO-associated spikes in the BLA could transfer information from the BLA to the cortex, like hippocampal sharp wave ripples (SPW-Rs)^31–34^. We therefore examined whether the BLA-HFOs modulate inter-regionally coordinated reactivations. The strength of assembly activations around the HFO onset was significantly enhanced during the first 30 min of post-NREM (0-30 min) compared with that of pre-NREM for with female condition, but not for without female condition (Fig. 4). Consistent with the time course of the increased reactivation rate during post-NREM sleep, the enhanced reactivation around the HFO onset was not observed in the second 30 min of post-NREM sleep (30-60 min) (Extended Data Fig. 9). Since the HFO-associated spiking of BLA neurons was not significantly different between pre-NREM and post-NREM in both conditions (Extended Data Fig. 8e-g), the enhancement of HFO-associated reactivations reflects inter-regional coordination rather than the sole modulation of BLA neuron spiking upon female presentation. Together, these results suggest that BLA-cortical inputs during BLA-HFOs act as a gating signal to selectively enhance the consolidation of a perceptual memory associated with emotional information during early NREM sleep by regulating inter-regional reactivations.

## NREM-specific BLA-cortical inputs enhance memory

Finally, we determined whether the post-NREM-specific BLA-cortical inputs causally contribute to the enhanced retention of texture memory using sleep-state-specific closed-loop optogenetics^17^ during the resting period of the associative learning task (Fig. 5a). We expressed an inhibitory opsin Jaws in the BLA and silenced BLA terminals at the M2 during the first 30 min of post-NREM period, then conducted recall sessions either on day 2 or 5 (Fig. 5b). This NREM-specific optogenetic manipulation did not alter sleep architecture (Extended Data Fig. 10a). On the day 2 recall sessions, unlike the chemogenetic experiments, both GFP-expressing control mice and Jaws-expressing mice showed a preference for the familiar side, and the total contact time did not differ between the two groups (Fig. 5c), indicating this manipulation did not suppress the associative consolidation of the memory for texture and female information. Next, we examined whether this manipulation affects texture memory enhancement tested during recall sessions on day 5. The GFP-expressing mice showed a preference for the familiar side, while the Jaws-expressing mice did not show the texture recognition, but there was still no difference in the total contact time (Fig. 5d). Next, we silenced BLA-M2 inputs during the post-REM period over a 10-hour time window and conducted recall sessions on day 5 (Fig. 5e). This REM-specific optogenetic manipulation did not alter sleep architecture (Extended Data Fig. 10b; total duration of post-REM sleep illumination in this 10-hour time window; GFP: 48.56 ± 2.89 min, Jaws: 46.04 ± 2.34 min). Unlike the post-NREM-specific silencing, both GFP-expressing and Jaws-expressing mice showed a preference for the familiar side and the total contact time did not differ (Fig. 5f). These results demonstrate that although BLA-M2 inputs during post-NREM do not contribute to the texture-female associative memory consolidation, these inputs are still necessary for the enhancement of the texture memory consolidation, and that BLA-M2 inputs during post-REM time periods are not causally related to the enhancement of the texture memory consolidation.

## Discussion

It has been hypothesised that the BLA modulates other brain regions to enhance a memory by emotional arousal, even if that memory is not stored in the BLA itself^2, 4^. Here, using the behavioural tasks which allowed us to examine the effect of emotional association on a neutral perceptual memory, we dissociated memory storage and consolidation from memory modulation. To the best of our knowledge, this provides the first behavioural evidence of the enhancement of the perceptual memory by emotional association in rodents, and we demonstrated the causal role of direct BLA-M2 inputs during NREM sleep in this perceptual memory enhancement (Fig. 5g).

The behavioural results showed that different aspects of associative memory (i.e., perceptual memory, emotional memory, and enhancement of perceptual memory) can be observed in a combination of different behavioural expressions (Fig. 1). Our results further clarify the role of the BLA and its projections to the dorsal frontal cortex in associative memory processing. Chemogenetic inactivation of M2-projecting BLA neurons affects all cellular processes, and this manipulation disrupted not only female memory consolidation but also texture memory enhancement (Fig. 2k-n). Furthermore, optogenetic silencing of BLA terminals on M2 during NREM sleep specifically disrupted the enhancement of texture memory (Fig. 5b-d). This difference suggests that female memory is consolidated in the BLA and/or its downstream areas (e.g., striatum, orbitofrontal cortex^4, 35^), but not in M2, where texture memory and its enhancement are coordinated. Although an alternative possibility is the difference in awake/sleep-state during the chemogenetic inactivation, a previous study has reported that even if PFC-projecting BLA neurons encode valence-specific information and are required for reward or punishment-based learning, BLA-PFC inputs do not trigger the valence-specific behaviour^24^, supporting our first assumption.

We identified M2 as a hub connecting the BLA and S1 (Fig. 2). Previous research has shown that plasticity in primary sensory cortices is crucial for the behavioural expression of memories that require perceptual recognition^17, 36^. Recent experimental evidence suggests that cortico-cortical^17, 37, 38^, thalamocortical^39, 40^, and neuromodulatory^41^ inputs convey the behaviourally relevant information and guide plasticity in sensory cortices^42^. Additionally, we have previously shown that M2-S1 top-down inputs generate dendritic calcium spikes followed by burst spiking in S1 neurons^43^, which is critical for the induction of plasticity^44^. This supports a recent theoretical study suggesting that burst-dependent plasticity precisely determines the loci of behaviourally relevant synapses according to the top-down inputs to update their weight in a hierarchical multi-layer network^45^. Thus, our findings propose a new circuit model for the memory regulation by emotion, in which the BLA interacts with top-down inputs and drives plasticity in lower-order cortical regions to enhance memory consolidation. Electrophysiological recordings showed that emotional association increased coordinated reactivation rates of learning-related activity across the BLA-M2-S1 regions during the first tens of minutes of NREM sleep, but not during REM sleep (Fig. 3). This time course of increased coordinated reactivations of the BLA and its associated regions (i.e., Hippocampus, PFC) is consistent with other studies using negative experiences^8, 9^. This suggests a general importance of reactivation across the BLA and other brain regions during early NREM sleep for emotional memory processing, regardless of emotional valence.

Notably, our data suggest that HFOs generated in the BLA^9, 29, 30^ act as an internal trigger to produce coordinated reactivations during early NREM sleep (Fig. 4). Previous studies have reported that SPW-Rs are generated locally in the hippocampus and trigger brain-wide neural activity during quiescence or NREM sleep^34^. Although the function of SPW-Rs in memory consolidation of the hippocampal and hippocampo-cortical systems has been elucidated^32, 33, 46, 47^, the function of BLA-HFOs has been unclear. Our findings will bring attention to BLA-HFOs and suggest that they play a role in modulating memory consolidation through direct BLA projections to other brain regions. In support of this idea, our optogenetic approach demonstrated the causal role of BLA-M2 inputs during this early NREM sleep for memory enhancement (Fig. 5). Elucidating the physiological roles of BLA-HFOs as well as hippocampal SPW-Rs is an intriguing topic for future research.

Finally, although we did not find a causal role for REM sleep-specific BLA-M2 inputs in the memory consolidation and enhancement in our task (Fig. 5e,f), other studies have shown the importance of REM sleep in hippocampus-dependent memory consolidation^48^, forgetting^49^, and thalamo-cortical circuit-dependent memory consolidation^50^. Altogether, our results highlight the importance of future research to investigate how circuit-dependent dynamics during a specific sleep-state contribute to different aspects of memory processing.

## Methods

### Animals

All animal experiments were performed under institutional guidelines and were approved by the Animal Experiment Committee of the RIKEN. Adult sexually naïve male and female wild-type mice (C57BL/6JJmsSlc, Japan SLC) older than 8 weeks were used in experiments. Mice were housed in a 12-h light/dark cycle in individually ventilated cages. All experiments used the male mice as experimental subjects, and female mice were used as natural reward for the male mice.

### General behavioural protocol

Before the behavioural tasks, we performed 3 days of handling and 3 days of habituation (30 min) in a black-walled open field arena (25 × 25 × 30 cm) with home cage bedding on the floor. Learning and recall sessions were conducted in the arena (25 × 25 × 30 cm) with two transparent dividing walls in the centre and two wired cups in opposite corners. The arena was black-walled to remove visual cues for the mice to distinguish between the two sides of the arena. The learning and recall sessions took place at the beginning of the light period. Mice were separated into two groups according to their resting periods. To test the memory consolidation, we provided a 24-h resting period and subsequently performed the recall sessions on day 2. For testing the enhanced retention of the texture memory, we used a 96-h resting period and performed recall sessions on day 5. Retention intervals were determined according to a previous study^17^. The arena was cleaned with 70% ethanol before each session and all behavioural apparatus was washed after each session. Texture memory was evaluated according to the preference for exploratory behaviour determined by the difference of cup contact time on the familiar texture floor (*T_familiar_*) and the novel texture floor (*T_novel_*): *Preference* (%) = (*T_familiar_* – *T_novel_*)/(*T_familiar_* + *T_novel_*) × 100. A positive value indicated that they preferred to explore the cup on the familiar texture floor side, and a negative value indicated that they preferred to explore the cup on the novel texture floor side. For the same texture control, preference was calculated by taking the cup contact time on the female-presented side minus the cup contact time on the empty side as the numerator and dividing by the sum of these values. Mice were divided into the following task conditions, and each mouse experienced only one learning and recall session.

### Perceptual recognition task

The perceptual memory task was conducted as described previously^17^. Briefly, we conducted this task as follows: On day 1, male mice were exposed to the arena containing the smooth floor for 30 min (learning session) and were then transferred to their home cage for resting. After a 24-h or a 96-h interval, texture memory retention was tested. Thus, on days 2 or 5, the mice were exposed to the arena containing the familiar smooth and a novel grooved textured (Batten 3.2-mm spacing) floors for 10- min. Mice do not have an innate preference for these smooth or grooved textures^17^.

### Associative recognition task

The associative memory task was conducted as follows: On day 1, male mice were exposed to the arena containing a female mouse in one of two cups on smooth floors. The mice were then transferred to their home cage for the 24-h or 96-h resting period prior to testing the texture memory retention. On days 2 or 5, mice were exposed to the arena containing both the familiar smooth and novel grooved texture floors for 10 min. Before the learning session, we measured the vaginal impedances (MK-11, Muromachi Kikai) of female mice to monitor their oestrus cycles. We then used a female mouse with vaginal impedance greater than 3 kΩ (oestrus state) as stimulus during the learning sessions.

### Associative recognition task: same texture control

To exclude the possibility that the mice used other memory, such as visual or spatial memory, rather than texture memory to guide their behaviour during the recall sessions, we performed the same texture control. In this task, the learning sessions were conducted exactly as that during the associative memory task. After a 24-h resting period, recall sessions were conducted in an arena containing only smooth floor on both sides.

### Virus vector preparation

DNA plasmids encoding pENN-AAV-hSyn-Cre-hGH (#105553) and pAAV-CAG-H2B-GFP (#116869) were obtained from Addgene. AAV production was performed as previously described^51^: AAV1-hSyn-Cre-hGH and AAVrg-CAG-H2B-GFP. The following AAVs were obtained from Addgene: AAV1-hSyn-EGFP (#50465-AAV1), AAVrg-CAG-FLEX-tdTomato (#28306-AAVrg), AAVrg-pgk-Cre (#24593-AAVrg), AAV1-EF1a-DIO-EYFP (#27056-AAV1), and AAV8-hSyn-DIO-hM4D(Gi)-mCherry (#44362-AAV8). AAV1-hSyn-Jaws-KGC-GFP-ER2 was obtained from UNC vector core.

### AAV injection

The mice were anesthetised using isoflurane. The scalp was shaved, cleaned with 70% ethanol and iodine, and then incised to expose the skull surface. The skull surface was levelled in the anterior-posterior direction by measuring the difference in the z-plane of the bregma and lambda. Then, the skull was levelled in the medio-lateral direction by moving the drill bit to −1.5 mm posterior to the bregma and measuring the difference in the z-plane ±1.5 mm from the midline. A craniotomy was performed over the injection sites. AAV vectors were injected using a pulled fine-glass capillary. Four weeks after AAV injection, the mice were either fixed for tissue collection or subjected to behavioural experiments.

### General protocol for histological experiments

Mice were deeply anesthetised by intraperitoneal injection of urethane and perfused transcardially with Hanks’ balanced salt solution (14025-076, Gibco) supplemented with heparin (10 units/mL), followed by 4% paraformaldehyde (PFA) in phosphate buffer saline (PBS; pH 7.4). The brains were then harvested and post-fixated in PFA overnight at 4 °C. The brains were sliced into either 40-μm or 50-μm coronal sections. For 40-μm sections, the brains were cryoprotected with 30% sucrose in PBS and sliced using a freezing microtome (ROM-380, Yamato). Meanwhile, for 50-μm sections, the brains were embedded in 4% agar (01059-85, Nacalai Tesque) and sliced using a vibratome (VT1200S, Leica). For immunohistochemistry, the sections were incubated in a blocking solution (5% normal goat serum and 0.3% TritonX-100 in PBS) for 30 min at room temperature. The sections were then incubated in a blocking solution containing a primary antibody, overnight at 4 °C, and subsequently washed with a washing buffer (0.3% TritonX-100 in PBS) three times. The sections were incubated in a blocking solution containing a secondary antibody conjugated with a fluorescent dye for 2 h at room temperature and washed three times with the washing buffer. The sections were mounted onto slides using a Fluoromount/Plus anti-fading agent (K048; Diagnostic BioSystems).

### Quantification of c-Fos+ cells

Male mice were handled and habituated in accordance with the aforementioned behavioural protocol. One day before the learning sessions, the mice were single housed. The mice were exposed to the behavioural arena with or without a female for 30 min and then transferred to their home cage. Two hours after the learning session, the mice were fixed. The brains were sliced into 50-μm sections, and immunohistochemistry was performed. We used an anti-c-Fos antibody (rabbit, 1/1000, #SC-52, Santa Cruz Biotechnology) as the primary antibody and anti-rabbit IgG antibody conjugated with Alexa Fluor 488 (goat, 1/400, #A11034, Molecular Probes) as a secondary antibody. Counterstaining was performed with 4′,6-diamidino-2-phenylindole (DAPI) (D9542, Sigma) during incubation with the secondary antibody. Five sections at 100-μm intervals containing representative BLA coordinates for each mouse were imaged using an inverted microscope (IX83, Olympus) equipped with a ×10 objective (UPLSAPO 2 10x / 0.40, Olympus). To calculate the density of c-fos+ cells in the BLA, the total c-Fos+ cells across the five sections was counted and divided by the total area of the BLA.

### AAVs and injection coordinates

All AAVs were injected into the structures of the right hemisphere. To trace BLA axons, we injected 150 nl of AAV1-hSyn-EGFP (titre 1.1 × 10^12^ GC/ml) into the BLA (AP, −1.4 mm, ML, 3.33 mm and DV, −4.2 mm). To trace M2-projecting neurons, we injected 300 nl of AAVrg-CAG-H2B-GFP (titre 2.38 × 10^12^ GC/ml) into the M2 (AP, 2.19 mm, ML, 0.6 mm, DV, 50 nl in −0.6 mm, 100 nl in −0.3 mm, and 150 nl in −0.1 mm). To trace BLA-recipient S1-projecting neurons, we injected with 150 nl of AAV1-hSyn-Cre-hGH (titre 1.7 × 10^13^ GC/ml) in the BLA (AP, −1.4 mm, ML, 3.33 mm and DV, −4.2 mm) and 300 nl of AAVrg-CAG-FLEX-tdTomato (titre 7.5 × 10^12^ GC/ml) into the S1 (AP, −0.73 mm, ML, 1.95 mm, DV, 100 nl in −0.6 mm, 100 nl in −0.3 mm, and 100 nl in −0.1 mm). To trace M2-projecting BLA axons, we injected with 300 nl of AAVrg-pgk-Cre (titre 1 × 10^13^ GC/ml) into the M2 (AP, 2.19 mm, ML, 0.6 mm, DV, 50 nl in −0.6 mm, 100 nl in −0.3 mm, and 150 nl in −0.1 mm) and 100 nl of AAV1-EF1a-DIO-EYFP (titre 2.6 × 10^12^ GC/ml) into the BLA (AP, −1.4 mm, ML, 3.33 mm, and DV, −4.2 mm).

### Immunohistochemistry

The fixed brains were sliced into 40-μm sections. To trace BLA axons and M2-projecting BLA axons, we performed immunohistochemistry using an anti-GFP antibody (rabbit, 1/2000, #598, MBL) as the primary antibody and an anti-rabbit IgG antibody conjugated with Alexa Fluor 488 plus (goat, 1/400, #A32731, Thermo Fisher Scientific) as a secondary antibody. Counterstaining was performed with NeuroTrace 435/455 (#N21479, Thermo Fisher Scientific) or NeuroTrace 640/660 (#N21483, Thermo Fisher Scientific) simultaneously with incubation with the secondary antibody. Images were acquired using an inverted microscope (IX83, Olympus) equipped with a ×4 objective (UPLSAPO 4x / 0.16, Olympus) or a confocal laser scanning microscope (FV3000RS, Olympus) with a ×10 objective (UPLSAPO 2 10x / 0.40, Olympus). To trace BLA-recipient S1-projecting neurons, sections were incubated with NeuroTrace 640/660 (#N21483, Thermo Fisher Scientific) for counterstaining. To confirm the injection site of AAV1-hSyn-Cre-hGH, we performed immunohistochemistry using an anti-Cre antibody (mouse, 1/1000, #MAB3120, Millipore) as a primary antibody and an anti-mouse IgG antibody conjugated with Alexa Fluor 488 (goat, 1/400, #A11029, Molecular Probes) as a secondary antibody. To trace M2-projecting neurons, sections were incubated with NeuroTrace 640/660 (#N21483, Thermo Fisher Scientific) for counterstaining. Images were acquired using an inverted microscope (IX83, Olympus) equipped with a ×4 objective (UPLSAPO 4x / 0.16, Olympus) or a confocal laser scanning microscope (FV3000RS, Olympus) with a ×20 objective (UPLSAPO 20X 20x / 0.75, Olympus).

### Quantification of *Fezf2*+ cells

To examine the expression of *Fezf2* mRNA in M2-projecting BLA neurons, we used the RNAscope Multiplex Fluorescent Reagent Kit v2 assay (Advanced Cell Diagnostics). Brain sections labelled for M2-projecting neurons with H2B-GFP, which contained representative BLA coordinates, were used. Sections were washed four times with 1× Tris-buffered saline (TBS) and incubated with RNAscope hydrogen peroxide for 30 min at room temperatures. After washing twice with 1×TBS and twice with 0.5×TBS, sections were mounted on Superfrost Plus slides (Thermo Fisher Scientific) and baked in a dry oven for 1 h at 60 °C. Then, sections were boiled in 1x Target Retrieval Reagent for 15 min at 90 °C and washed with distilled H_2_O and 100% ethanol, after which they were baked in a drying oven for 30 min at 60 °C. Sections were incubated with RNA scope Protease Plus in a dry oven for 30 min at 40 °C and washed twice with distilled H_2_O. Then, sections were incubated with a probe against *Fezf2* (#888591-C2, Mm-Fezf2-O2-C2) for 2 h at 40 °C, and further, amplifying hybridisation processes were performed with AMP1, AMP2, and AMP3 sequentially. Horseradish peroxidase (HRP) signal processing was performed using HRP-C2 and Opal570. Subsequently, immunohistochemistry was performed using an anti-GFP antibody (chicken, 1/400, #ab13970, Abcam) as the primary antibody, an anti-chicken IgY antibody conjugated with Alexa Fluor 488 plus (goat, 1/200, #A32931, Thermo Fisher Scientific) as the secondary antibody, and NeuroTrace 640/660 (#N21483, Thermo Fisher Scientific) for counterstaining. Images were acquired using either an inverted microscope (IX83, Olympus) equipped with a ×4 objective (UPLSAPO 4x / 0.16, Olympus) or a confocal laser scanning microscope (FV3000RS, Olympus) with a ×20 objective (UPLSAPO 20X 20x / 0.75). Images were processed using the ImageJ/Fiji software. We stacked the images with max z-projection, and the background of the Fezf2 channel was subtracted using a rolling ball and median filter. We then counted *Fezf2*+ and *Fezf2*- neurons in GFP+ M2-projecting neurons.

### Anatomical reconstruction

We used BrainJ (https://github.com/lahammond/BrainJ) in ImageJ/Fiji to reconstruct the M2- projecting BLA axons. Sections were imaged at 40-μm intervals using an inverted microscope (IX83, Olympus) equipped with a ×4 objective (UPLSAPO 4x / 0.16, Olympus). Images were processed and registered in the Allen Brain Atlas Common Coordinate Framework with BrainJ, as described previously^52^. Visualisation of the 3D reconstruction was performed using the Volume Viewer plugin in ImageJ/Fiji.

### Chemogenetic experiment

For expression of hM4Di in M2-projecting BLA neurons, we injected with 300 nl of AAV8-DIO-hM4Di (titre 4.9 × 10^12^ GC/ml) in the bilateral BLA (AP, −1.4 mm, ML, ±3.33 mm and DV, −4.2 mm) and 300 nl of AAVrg-pgk-Cre (titre 9.5 × 10^12^ GC/ml) in the bilateral M2 (AP, 2.19 mm, ML, 0.6 mm, DV, 50 nl in −0.6 mm, 100 nl in −0.3 mm, 150 nl in −0.1 mm). Four weeks after AAV injections, mice were handled for 3 days and habituated to the behavioural arena for 3 days, then the associative memory task was conducted. Immediately after the learning sessions, saline or clozapine N-oxide (CNO) (1 mg/kg, intraperitoneally) were administered, and the mice were returned to the home cage. Recall sessions were conducted on day 2 or day 5 to test texture-female associative memory consolidation and texture memory enhancement, respectively.

### Polysomnographic recording

Electroencephalography (EEG) and electromyography (EMG) recordings were performed as previously described ^17^. Briefly, we implanted an EEG screw (M0.8 × 2 mm, TE-00013, Matsumoto) in the parietal cortex and a reference screw in the cerebellum for EEG recording and two flexible wires (AS 633, Cooner Wire) in the neck muscle for EMG recording. Dental cement was used to cover the screws and attach a connector to the skull. EEG signals from the parietal cortex were referenced to the cerebellum screw. For habituation, mice were connected to recording wires at least 3 days prior to recording. Continuous EEG and EMG recordings were performed after the learning sessions of the perceptual (without female) or the associative (with female) memory tasks. EEG and EMG signals were recorded at a sampling rate of 10 kHz using an NI-DAQ (NI-9215, National Instruments) with a custom-made LabVIEW software. Sleep scoring was performed as described in a previous report^17^. Briefly, the root mean square (RMS) of EMG signals, EEG delta (1-4Hz) power (normalised to power in 1-50 Hz) and theta (6-9 Hz) power (normalised to power in 1-4 Hz) were calculated in each 4-s sliding window. We defined brain states as follows; awake− RMS value of EMG was above the threshold which was defined manually in each animal; NREM− RMS value of EMG was below the threshold and delta power/theta power was above 0.1. REM− RMS value of EMG was below the threshold and delta power/theta power was below 0.1. When the tentative state of a window, which differs from the current state, continued through three consecutive windows, the current state changed to a state that was tentatively defined by the data from the three epochs. The state transition from awake to REM was denied.

### Single unit recording

A custom-designed microdrive (https://github.com/yoshihito-saito/microdrive) containing four tetrodes targeting the M2, five tetrodes targeting the S1, and two octrodes targeting the BLA was constructed. Each electrode was made of tungsten wire (12.5 μm, California Fine Wire), and typically had an impedance of 200-1000 kΩ at 1 kHz. The EEG screw, reference screw, and EMG wires were connected to an electrode interface board (EIB) (64-ch Hirose EIB, OpenEphys). Mice were anaesthetised using isoflurane and injected intraperitoneally with carprofen (5 mg/kg), baytril (10 mg/kg), and dexamethasone (4 mg/kg). The scalp was shaved, cleaned with 70% ethanol and iodine, and then incised to expose the skull surface. The skull surface was levelled in the anterior-posterior direction by measuring the difference in the z-plane of the bregma and lambda. Then, the skull was levelled in the medial-lateral direction by moving the drill bit −1.5 mm posterior to the bregma and measuring the difference in the z-plane ±1.5 mm from the midline. Craniotomy and duratomy were performed above the recording sites. An EEG and reference screw and EMG wires were implanted. The microdrive was subsequently implanted over the right hemisphere and the surface of the brain covered with a drop of silicon oil. The microdrive was fixed to the skull using dental cement. The position of each electrode was adjusted daily to reach the target depth. Four days after surgery, mice were transferred to a recording home cage with bedding and habituated for one week before recording. Mice were handled and habituated to the behavioural arena, and then, the perceptual (without female) or associative (with female) memory tasks were conducted. Neural activity was recorded as follows: 1 h in the recording home cage prior to, 30 min in the behavioural arena during, and 3 h in the recording home cage after the learning session. Signals were amplified and digitised with a 64-channel headstage (Low-profile SPI Headstage 64ch, OpenEphys) and recorded at 30 kHz using an Intan RHD USB interface board (IntanTech).

### Data processing

Extracellular signals were processed with a pipeline made by Spikeinterface (https://github.com/SpikeInterface/spikeinterface)^53^ as follows: (1) signals were filtered at 600-6000 Hz and applied a common median filter; (2) filtered signals were sorted with MoutainSort4; (3) sorted units were manually curated with Phy and units with inter-spike intervals violation < 0.5 were included; and (4) units were classified into regular spiking (RS) and fast spiking (FS) by their peak to valley width of the waveforms (RS > 0.4 ms, FS < 0.4 ms).

### State-dependent spike rate property analysis

Brain states were classified using simultaneously recorded EEG and EMG signals as described above. Spike rates during learning sessions, post-NREM and post-REM were calculated for each unit. The spike rate (*SR*) changes between states were calculated as (*State* 1 *SR* – *State* 2 *SR*)/ (*State* 1 *SR* + *State* 2 *SR*) for each state pair.

### Assembly detection and reactivation analysis

We detected coordinated activity patterns (neural assembly) as follows^27, 28, 54^: (1) spike trains of RS and FS were binned into 25-ms intervals; (2) Pearson correlation coefficients for all pairs of neuronal activity during a learning epoch were calculated, and the correlation matrix was created by concatenating all pairs of correlation coefficients; (3) principal component analysis (PCA) was applied to the matrix; (4) the number of neural assembly was determined as the number of eigenvalues exceeding the Marčenko–Pastur distribution threshold:

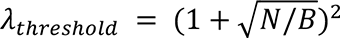

where *N* and *B* represent the number of units and time points, respectively; (5) independent component analysis (ICA) was applied to each assembly to compute the contribution of each unit to the assembly as the vector of weights (*N* length); (6) each weight vector *w* was normalised to unit length:

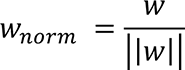

(7) units with the weight exceeding the threshold 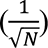 were defined as member units because if each unit composed equal contribution to the assembly, the weights of all units should be 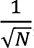; (8) assemblies which contain at least a single member unit in each region were defined as “tri-regional assembly” and used for further analysis.

We then tracked the assembly pattern reactivation strength during post-NREM and post-REM (target state) as follows: (1) a projection matrix *P* was constructed by calculating the outer product of its weight vector:

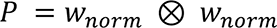

followed by the diagonal setting to zero; (2) reactivation strength was calculated as

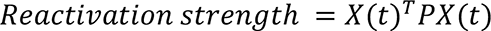

where *X*(*t*) represents the z-scored binned spike counts (25-ms, N-by-number of time points) during a target state at the given time point *t*. We defined a reactivation event as when the peak of the reactivation strength exceeded the threshold of 5.

### Cross-correlograms of inter-regional cell pairs during reactivations

To calculate the cross-correlogram between one member unit in region 1 (reference unit) and another member unit in region 2 (target unit) during reactivations, spike trains of member units were binned into 1-ms intervals. We then z-scored the target unit’s binned spikes. For each spike in a reference unit during each reactivation event, the z-scored binned spikes of the target unit were incremented and normalised by the number of reactivation events.

### BLA High-frequency oscillations

BLA-HFOs were detected based on the previously described method^9^. The median of the LFPs from each electrode in the BLA was down-sampled to 1000 Hz, band-pass filtered at 100–125 Hz, and Hilbert transformed to extract the instantaneous amplitude. Signals were then converted to z-score. Periods with a z-score > 2 were identified as candidate events, and candidates within a 20-ms window were concatenated into a single candidate. Candidates were classified as HFO if the duration of events was greater than 30 ms and the peak amplitude of events was greater than 3 z.

To analyse the assembly activation associated with BLA-HFOs, we z-scored the assembly strengths and calculated peri-event triggered averages in a ±1 s window.

### Optogenetic experiment

For optogenetic experiments, we injected with 200 nl of AAV1-hSyn-EGFP (titre 1.1 × 10^12^ GC/ml) or AAV1-hSyn-Jaws-GFP (titre 1.2 × 10^12^ GC/ml) in the bilateral BLA (AP, −1.4 mm, ML, ±3.33 mm and DV, −4.2 mm). The Teleopto system (Bio Research Center) was used for optogenetic experiments. Two weeks after the AAV injection, LED device implantation and polysomnographic recording surgery were performed as previously described^17^. Briefly, optic fibres (NA 0.39, Ø400 μm, FT400EMT, Thorlabs) were attached to each end of a doublet LED (590 nm, ∼1.5 mW at the fibre tip, TeleLP-Y-covered, Bio Research Center). The LED device, EEG screws and EMG wires were implanted on the brain surface of the bilateral M2, parietal cortex and neck muscle, respectively, and fixed to the skull with dental cement. One week after surgery, mice were transferred to a recording home cage with bedding and habituated for one week before initiating recording in their home cage. Mice were managed and habituated to the behavioural arena, and then the associative memory task was conducted. For sleep-state-specific photostimulation, EEG and EMG signals were recorded at a sampling rate of 10 kHz by a NI-DAQ (NI-9215, National Instruments) with a custom-made LabVIEW software and online sleep scoring was performed as described in above. Photostimulation was started from the onset of transition to a target state and lasting to transition to other states. For NREM-specific photostimulation, the total duration of photostimulation was pre-set to 30 min. Recall sessions were performed on day 2 or day 5 to test texture-female associative memory consolidation and texture memory enhancement, respectively. For REM-specific photostimulation, photostimulation was performed during the entire REM sleep period within the 10-h time window, and recall sessions were conducted on day 5 to test texture memory enhancement.

### Statistics and data presentation

All statistical tests used are indicated in the respective subsection. All statistical analyses were performed using Python, and all statistical tests were two-sided. Statistical significances are shown as n.s., *, **, and *** indicate *P* ≥ 0.05, *P* < 0.05, *P* < 0.01, and *P* < 0.001, respectively. For anatomical, chemogenetic, and optogenetic experiments, animals whose viral injection location had fallen outside the target region were excluded from the analysis. For electrophysiological experiments, animals were excluded if the electrode became stuck before reaching the target region. Units recorded from outside the target region were also excluded from analysis. For anatomical experiments, at least three mice were used, and the same trend was observed. Representative images are shown in the respective figures.

## Data availability

Data supporting the findings of this study are available from the corresponding author on reasonable request.

## Code availability

Custom analysis codes for electrophysiological experiments and the Microdrive design are available on https://github.com/yoshihito-saito.

## Acknowledgements

We thank Y. Atsumi, Y.Ito, H. Kazama, A. Noguchi, T. Ohnuki, A. Ohuchi, H. Okamoto, N. Ota, Y. Takema, T. Toyoizumi, K. Yamada, K. Yoshida, Y. Yoshihara and all lab members for comments and discussions on the project; A. Kamoshida for LabVIEW program development; R. Kato, Y. Mizukami and K. Ueno for animal care. This project was supported by Grant from Kao Corporation (to M. Murayama); AMED-Brain/Minds Project (JP15dm0207001 to M. Murayama); Grant-in-Aid for Transformative Research Areas (B) from the JSPS (20B305 to M. Murayama); Grant-in-Aid for Young Scientists (A) from the JSPS (16H05929 to M. Murayama); Junior Research Associate program of RIKEN (to Y.S.).

## Author contributions

Y.S., J.P.J., and M. Murayama conceptualized the project. Y.S. and M.O. conducted the experiments. Y.S. and Y. Osako analysed the data. Y. Oisi, C.M., S.K., and K.K. produced AAV vectors. M. Morita, and M. Murayama supervised the project. Y.S., J.P.J., and M. Murayama wrote the manuscript.

---

**Extended Data Fig. 1.**
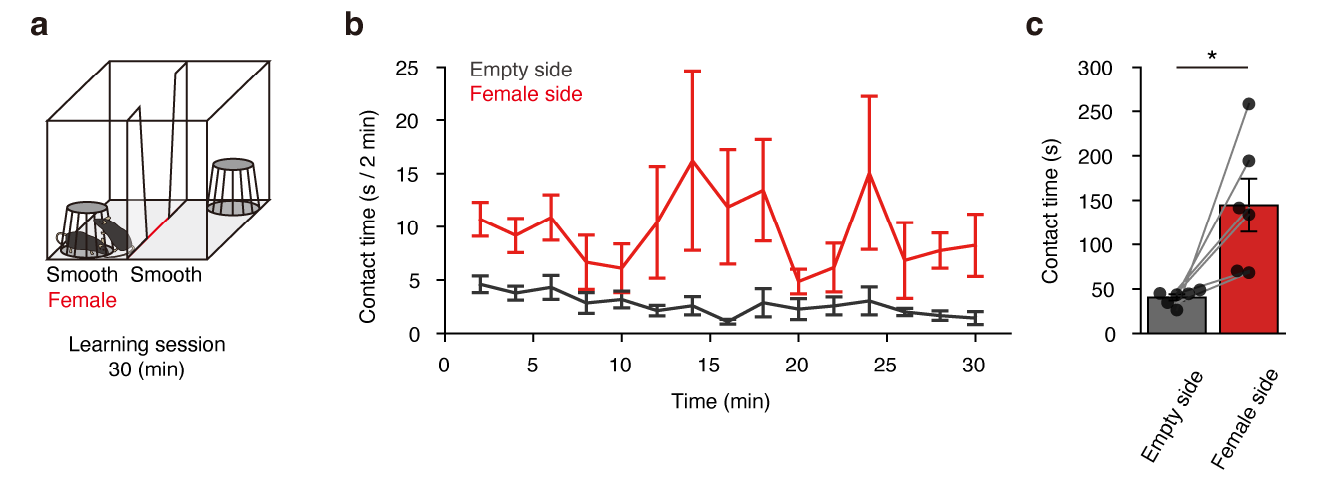
Male mice preferentially explore the female-presenting side of the arena. **a**, Schematic diagram of a learning session for the associative memory task. **b**, Time course of exploration behaviour during learning sessions of the associative memory task (*n* = 6). **c**, Total contact time for each side during the learning sessions (* *P* = 0.0312, Wilcoxon signed rank test). Data are presented as mean ± SEM.

**Extended Data Fig. 2.**
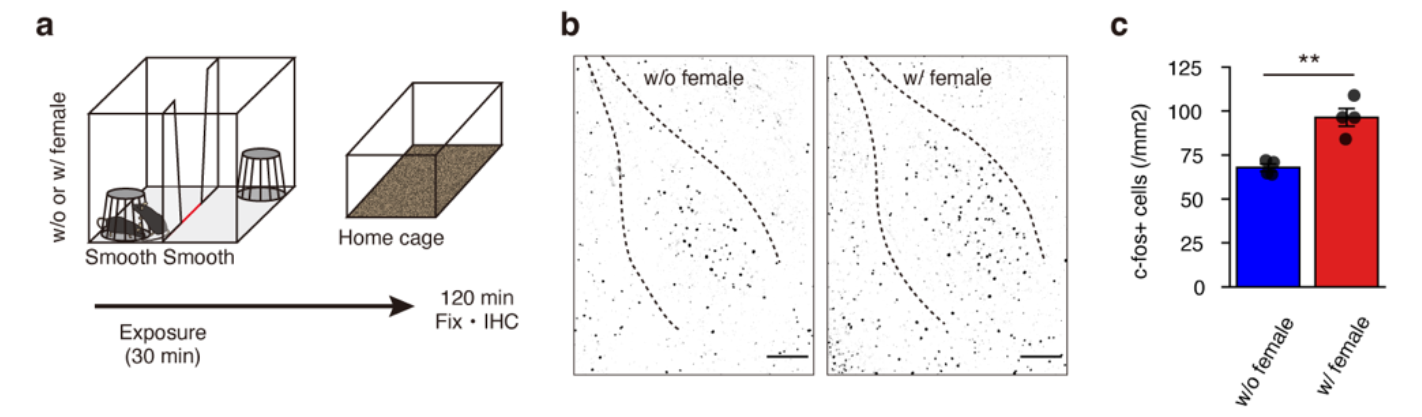
Female presentation in the arena increases c-Fos expression in the BLA. **a**, Schematic diagram of the c-Fos experiment. **b**, Example of the c-Fos immunostaining in the BLA after exposure to the arena without or with female presentation (scale bar, 200 μm). **c,** Quantification of the c-fos expression in the BLA (w/o female; *n* = 4, w/ female; *n* = 4, *t*_(6)_ = −5.21, ** *P* = 0.0020, Student’s *t*-test). Data are presented as mean ± SEM.

**Extended Data Fig. 3.**
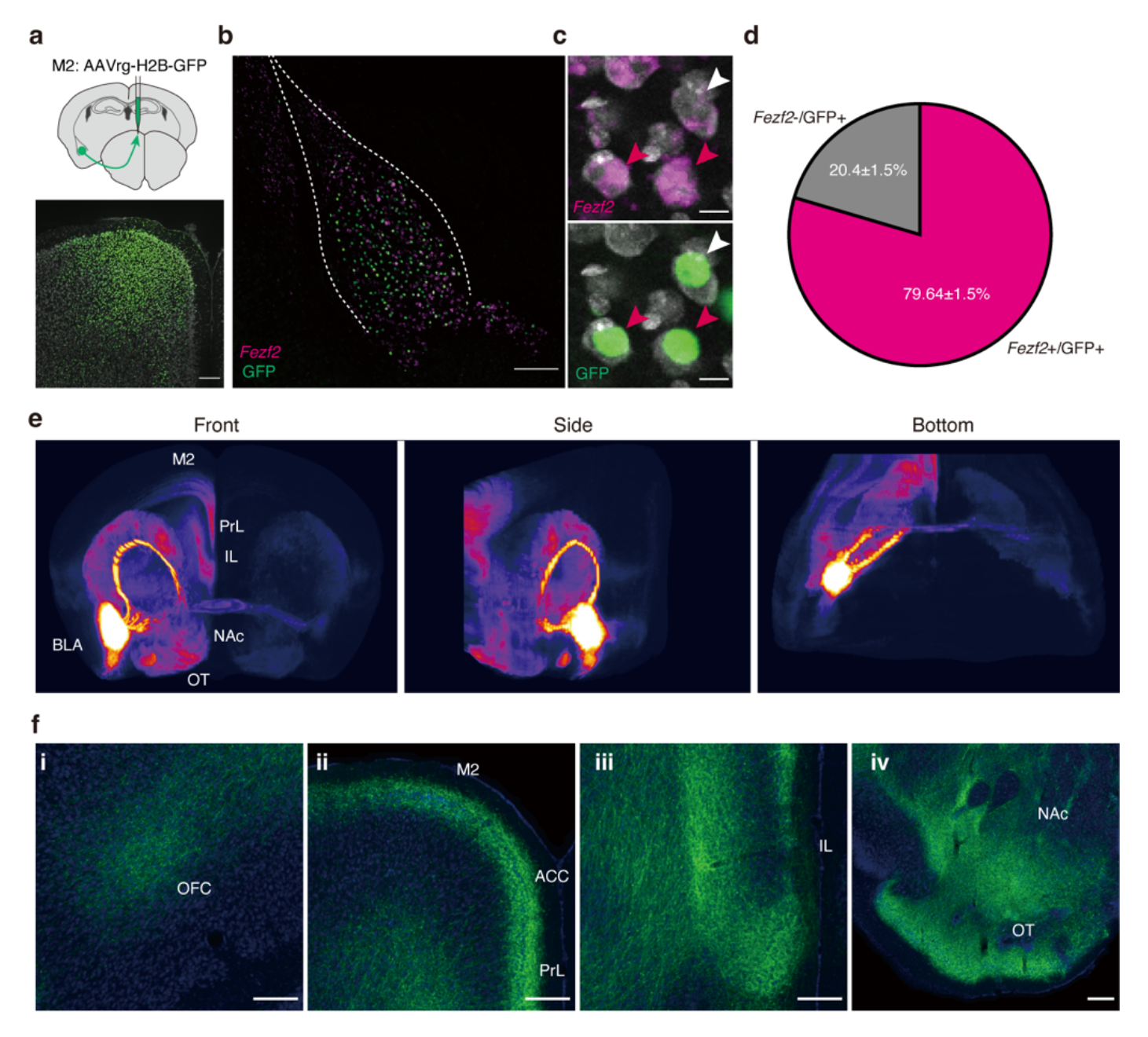
Molecular and anatomical features of M2-projecting BLA neurons. **a**, Retrograde labelling of M2-projecting neurons. AAVrg-H2B-GFP was unilaterally injected into the M2. **b**, *Fezf2* expression in M2-projecting BLA neurons. M2-projecting BLA neurons were labelled by injecting AAVrg-H2B-GFP in the M2, and *Fezf2* expression in the BLA was detected by RNAscope (scale bar, 200 μm). **c**, Magnified view. Red arrowheads indicate *Fezf2*+/H2B-GFP+ and white arrowheads indicate *Fezf2*-/H2B-GFP+ (scale bar, 10 μm). **d**, Quantification of the fraction of M2- projecting neurons expressing *Fezf2* (*n* = 3 mice, mean ± SEM). **e**, Brain wide distributions of M2- projectiong BLA axon collaterals labelled by injecting AAVrg-Cre and AAV-DIO-eYFP into the M2 and BLA, respectively. Fluorescence intensity of eYFP is presented as a heatmap. **f**, Magnified views of M2-projecting BLA axons in frontal cortical regions and striatal regions. (i) Orbitofrontal cortex, OFC, (ii) Secondary motor cortex, M2; Anterior cingulate cortex, ACC; Prelimbic cortex, PrL, (iii) Infralimbic cortex, IL, (iv) Olfactory tubercle, OT; Nucleus accumbens, NAc (scale bar, 200 μm).

**Extended Data Fig. 4.**
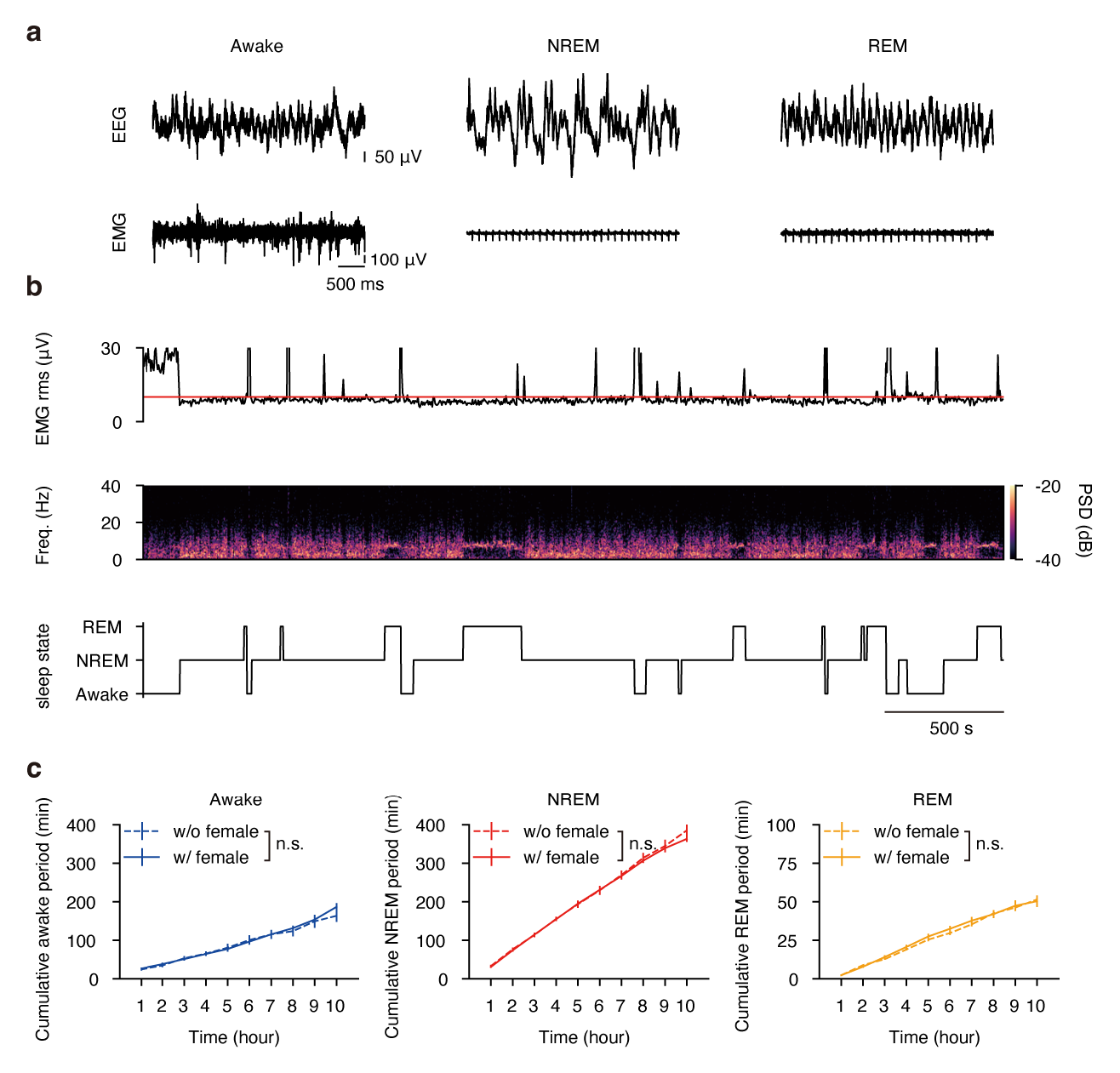
Female presentation in the arena does not alter sleep architecture. **a**, Sleep-state classification by simultaneous EEG and EMG recordings. Example EEG and EMG traces during awake (left), NREM (middle), and REM (right) states. **b**, Example of the post-learning sleep-state. Shown are EMG rms amplitude, EEG power spectrogram, and sleep states. **c**, Post-learning cumulative duration of awake, NREM and REM across 10 h in the with female and the without female groups (n.s. *P* > 0.05, Two-way repeated measures ANOVA). Data are presented as mean ± SEM.

**Extended Data Fig. 5.**
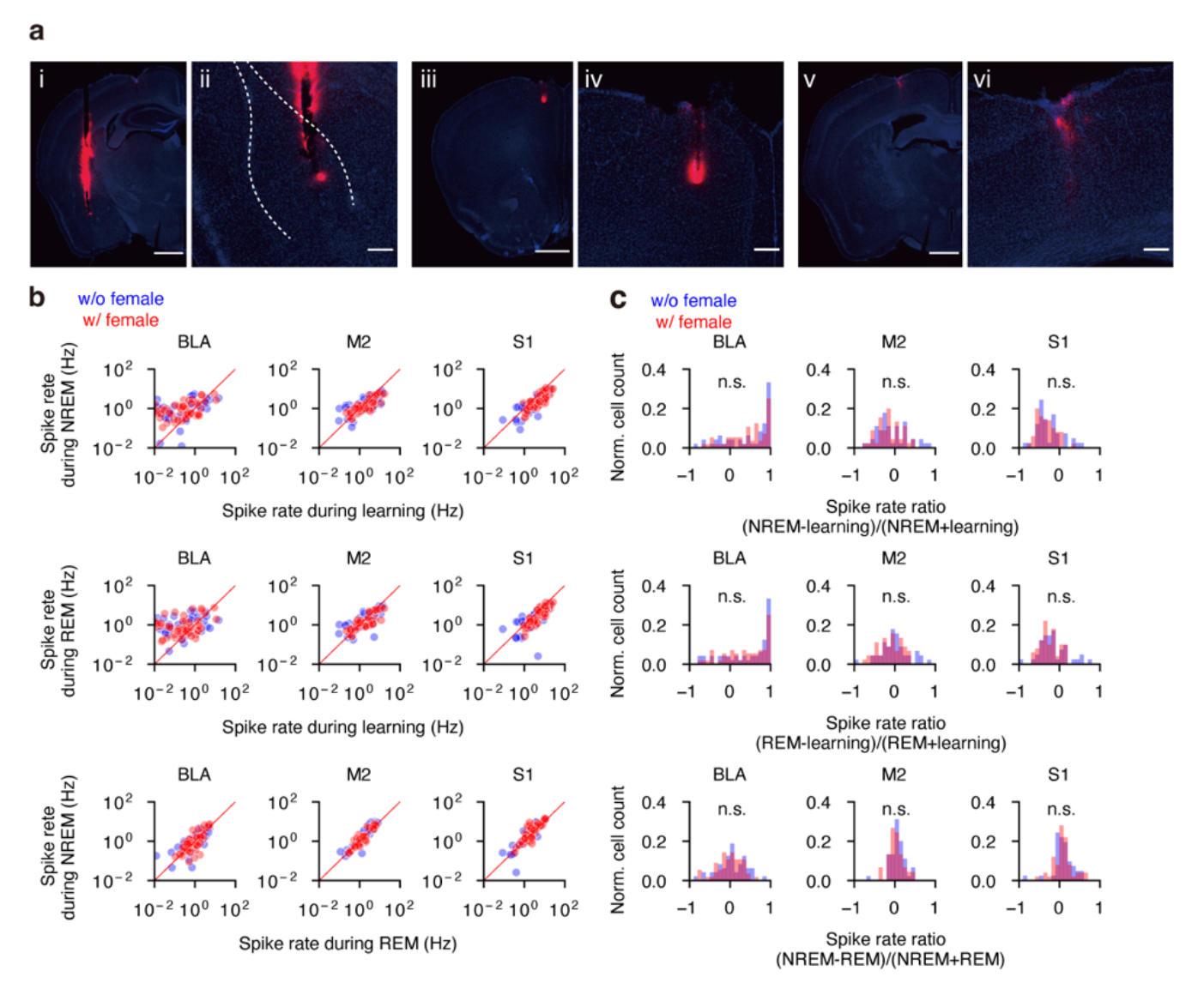
State-dependent spike rate changes of BLA, M2 and S1 units. **a**, Simultaneous recording from the BLA, the M2 and the S1. Images show examples of the electrode tracks in the BLA (i, ii), M2 (iii, iv), and S1 (v, vi) (scale bar: i, iii and v, 1 mm; ii, iv and vi, 200 μm). **b**, Sleep-state dependent spike rate shifts. Distributions of spike rates of units in the BLA, the M2 or the S1 during the learning session vs. post-NREM (top), learning session vs. post-REM (middle) and post-REM vs. post-NREM (bottom) in the with (red) and the without (grey) female groups. Each dot represents a unit. **c**, Distributions of spike rate ratios of each state pair in the with (red) and the without (grey) female groups (n.s. *P* > 0.05, Kolmogorov-Smirnov test, *n* = 56 units in the BLA, 45 units in M2, 50 units in S1 in the with female group, *n* = 48 units in the BLA, 45 units in M2, 41 units in S1 in the without female group).

**Extended Data Fig. 6.**
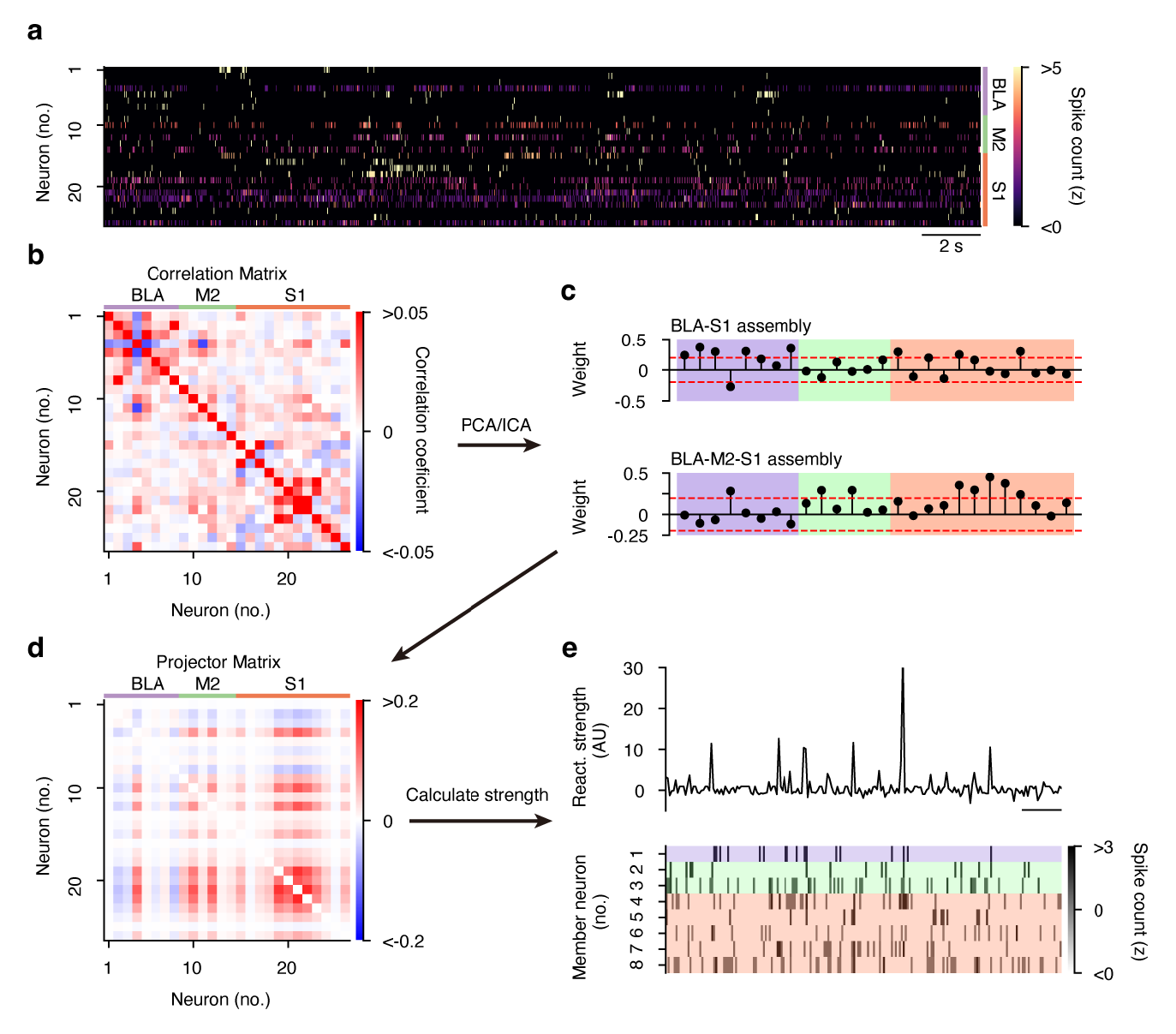
Detection of tri-regional assemblies. **a**, Example of the unit activity in the BLA (*n* = 8), M2 (*n* = 6), and S1 (*n* = 12) during a learning epoch in the with female condition. Spike trains were binned into 25 ms time bins and z-scored. **b**, Correlation matrix of the activity during the learning epoch. Principal component analysis (PCA) was applied to the correlation matrix to detect characteristic patterns. The number of significant assemblies was determined by the upper limit of the Marčenko–Pastur distribution. Independent component analysis (ICA) was applied to estimate the contribution of each unit to the detected assemblies. **c**, Two examples of the assembly patterns. A weight vector represents the contribution of each unit to an assembly. For each assembly, units with a high weight (exceeding the threshold (abs(1/√N), dashed red line) were defined as member units. Assemblies that contain at least a single member unit from each region were defined as “tri-regional assembly” (bottom) and used for further analysis. **d**, Projection matrix of the tri-regional assembly in (**c**). Projection matrix was constructed by calculating the outer product of the weight vector followed by the diagonal of the matrix set to zero. **e**, Example temporal reactivation of the assembly pattern (top) and z-scored binned spike counts of each member unit (bottom).

**Extended Data Fig. 7.**
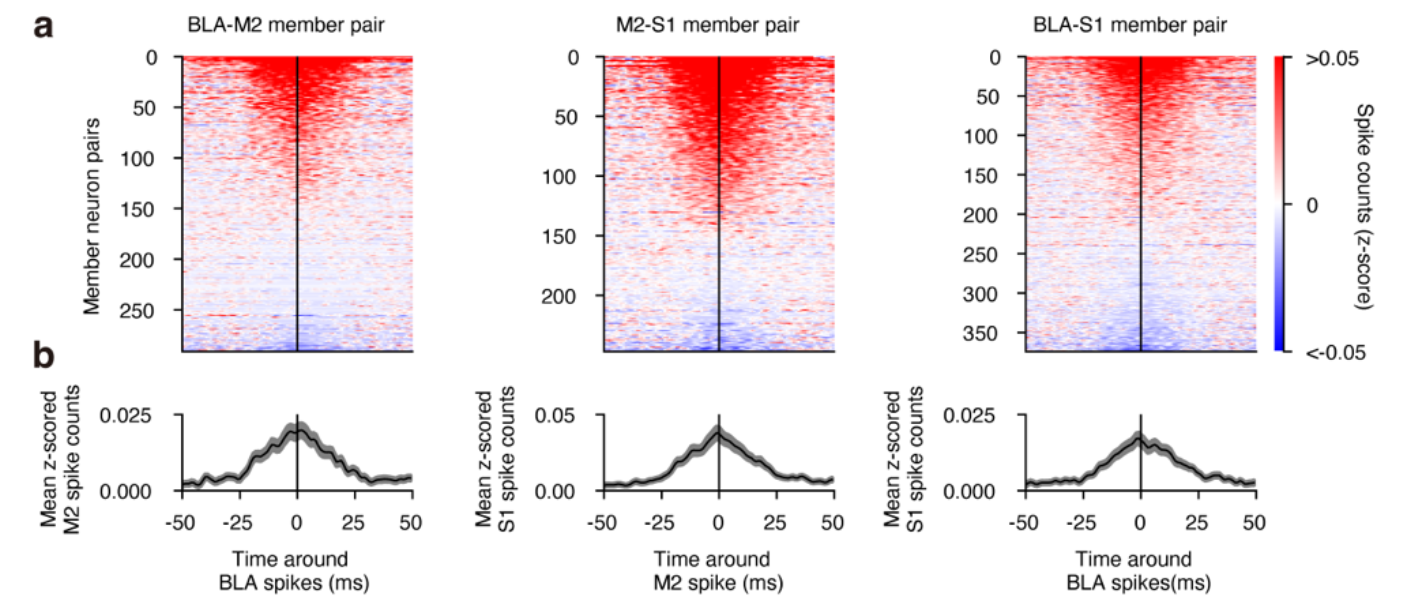
Inter-regional coordinated neural activity during reactivations. **a**, Cross-correlograms during reactivations of inter-regional member unit pairs from tri-regional assemblies. **b**, Mean (± SEM) cross-correlograms in (**a**).

**Extended Data Fig. 8.**
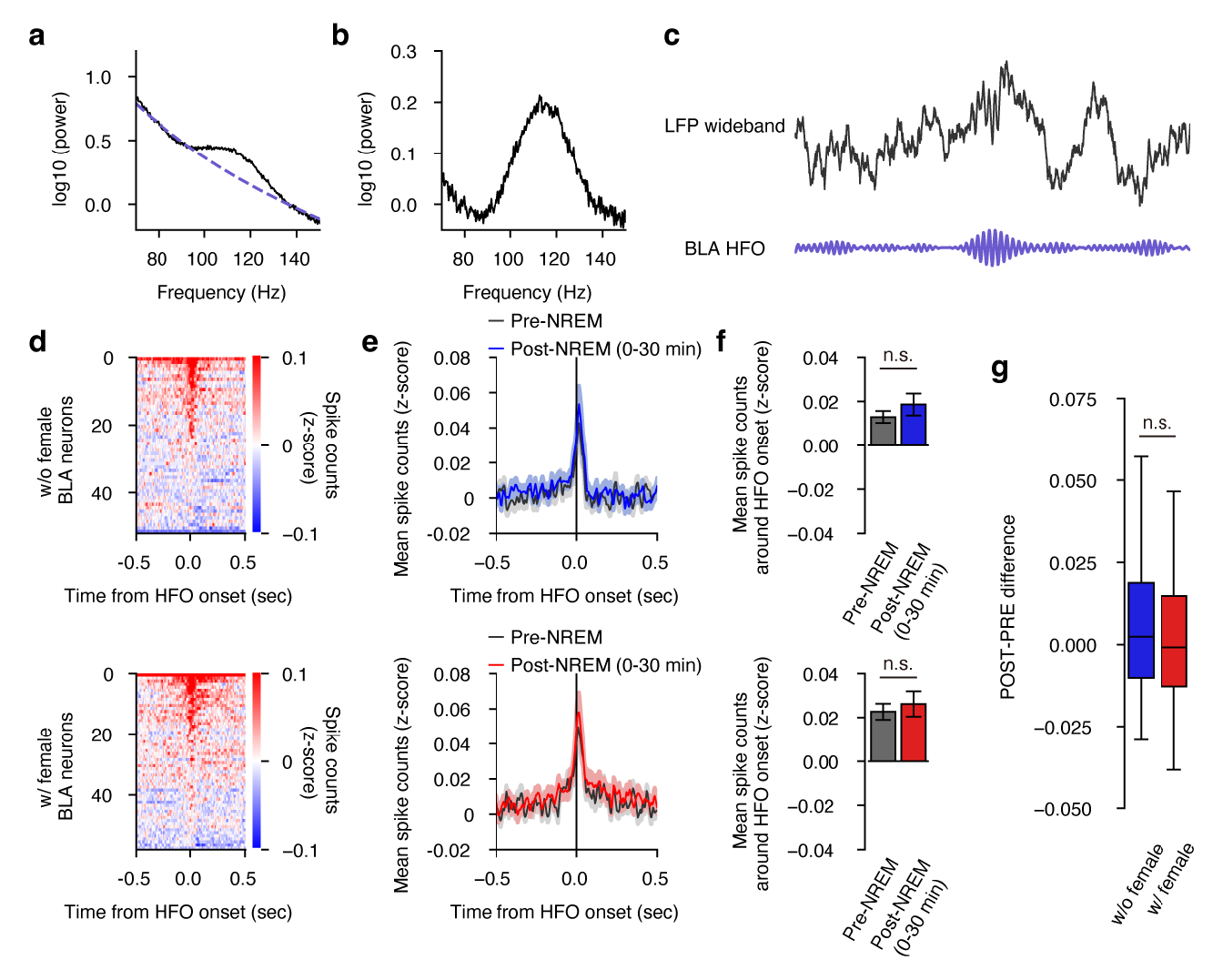
BLA-HFOs and associated spiking activity. **a**, An example of the power spectral density of local field potential (LFP) during NREM sleep in the BLA. The dashed line indicates the estimated aperiodic baseline of the signal. **b**, The baseline corrected power spectral density. Strong peak was observed in the 90-140 Hz range. **c**, An example of wideband and filtered LFP from the BLA. **d**, BLA-HFOs onset aligned firing of BLA units in without female condition (top) and with female condition (bottom). **e**, Mean (± SEM) z-scored spike counts of BLA units around BLA-HFOs onset in without female condition (top) and with female condition (bottom) during pre-NREM and first 30 min of post-NREM. **f**, Mean z-scored spike counts in a ± 100-ms window around BLA-HFOs onset in without female condition (top) and with female condition (bottom) during pre-NREM and post-NREM (0-30 min) (n.s. *P* > 0.05, Wilcoxon signed rank-test). **g**, The pre-NREM and post-NREM (0-30 min) difference in mean z-scored spike counts around the onset of BLA-HFOs was compared in the without female (blue) and with female (red) conditions (n.s. *P* > 0.05, Mann-Whitney U test). Boxplots show the median with 25/75 percentile (box) and 1.5 × interquartile range (whiskers).

**Extended Data Fig. 9.**
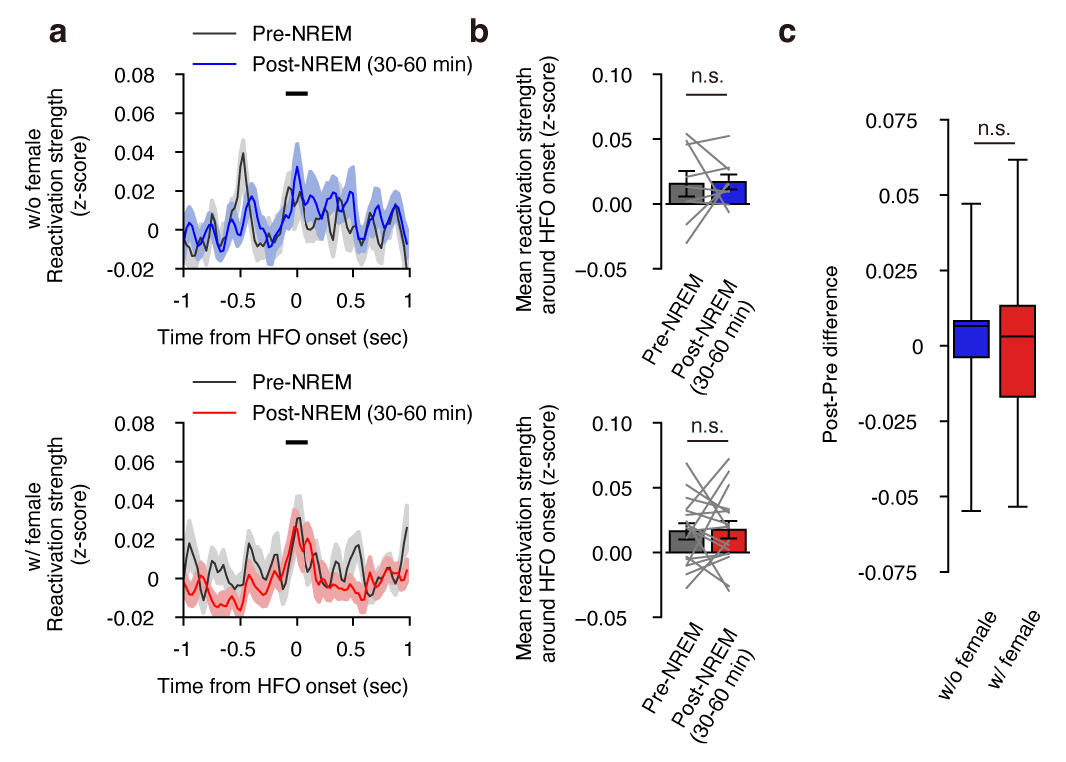
Inter-regional reactivation around BLA-HFOs during 30-60 min of NREM sleep. **a**, Mean z-scored reactivation strength of tri-regional assemblies around BLA-HFOs onset in without female condition (top) and with female condition (bottom) during pre-NREM and second 30-60 min of post-NREM (shade represents ± SEM). **b**, Mean reactivation strength in a ± 100- ms window around BLA-HFOs onset in without female condition (top) and with female condition (bottom) during pre-NREM and post-NREM (30-60 min) (without female, n.s. *P* = 0.6523; with female, n.s. *P* = 0.9632, Wilcoxon signed rank-test) **c**, The pre-NREM and post-NREM (30-60 min) difference in mean reactivation strength around the onset of BLA-HFOs was compared in the without female (blue) and with female (red) conditions (n.s. *P* = 0.3138, Mann-Whitney U test). Boxplots show the median with 25/75 percentile (box) and 1.5 × interquartile range (whiskers).

**Extended Data Fig. 10.**
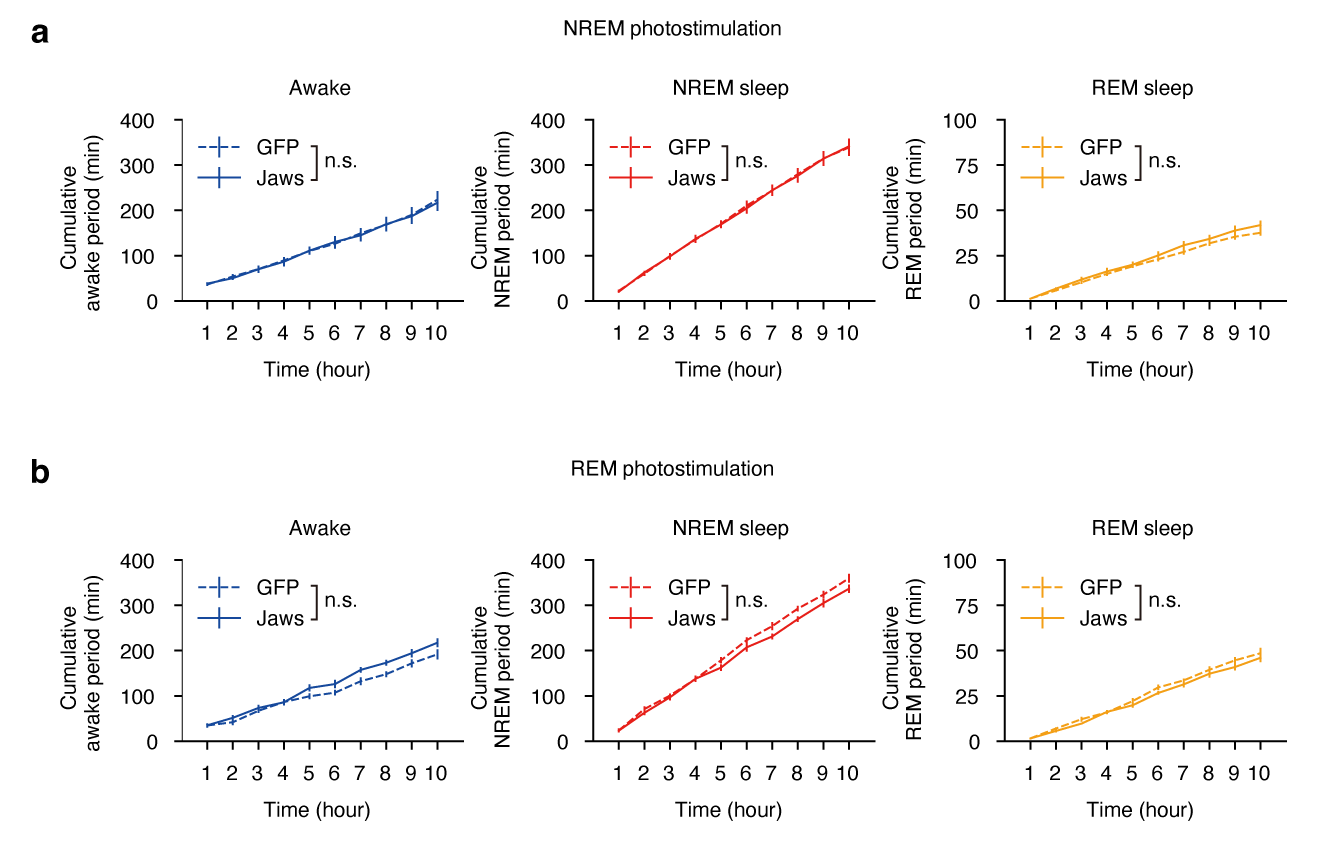
Optogenetic silencing does not alter sleep architecture. (**A**) Post-learning cumulative duration of being the awake, NREM, and REM states across 10-h in NREM-specific photostimulation groups (n.s. *P* > 0.05, two-way repeated measures ANOVA). (**B**) Post-learning cumulative duration of being awake, NREM and REM states across 10-h in REM specific photostimulation groups (n.s. *P* > 0.05, two-way repeated measures ANOVA). Data are presented as mean ± SEM.

**Supplementary Table 1.** Summary of recorded units and detected assemblies.

| MouseID | Condition | Number of units |  |  | Number of assemblies |  |  |  |  |  |  |
| --- | --- | --- | --- | --- | --- | --- | --- | --- | --- | --- | --- |
|  |  | BLA | M2 | S1 | BLA-M2-S1 | BLA-M2 | M2-S1 | BLA-S1 | BLA | M2 | S1 |
| TM015 | w/ female | RS: 25, FS: 0 | RS: 8, FS: 0 | RS: 10, FS: 1 | 8 | 0 | 0 | 0 | 0 | 0 | 1 |
| TM016 | w/o female | RS: 0, FS: 0 | RS: 11, FS: 0 | RS: 9, FS: 3 | 0 | 0 | 4 | 0 | 0 | 0 | 2 |
| TM018 | w/ female | RS: 0, FS: 0 | RS: 0, FS: 0 | RS: 1, FS: 0 | n.d. | n.d. | n.d. | n.d. | n.d. | n.d. | n.d. |
| TM019 | w/o female | RS: 14, FS: 2 | RS: 0, FS: 0 | RS: 1, FS: 4 | 0 | 0 | 3 | 1 | 0 | 0 | 0 |
| TM020 | w/ female | RS: 8, FS: 0 | RS: 2, FS: 0 | RS: 11, FS: 1 | 0 | 0 | 0 | 4 | 0 | 0 | 0 |
| TM022 | w/o female | RS: 0, FS: 0 | RS: 4, FS: 0 | RS: 7, FS: 0 | 0 | 0 | 0 | 0 | 0 | 0 | 2 |
| TM023 | w/o female | RS: 7, FS: 0 | RS: 0, FS: 0 | RS: 0, FS: 1 | 0 | 0 | 0 | 1 | 0 | 0 | 0 |
| TM025 | w/ female | RS: 6, FS: 0 | RS: 0, FS: 0 | RS: 0, FS: 0 | 0 | 0 | 0 | 0 | 1 | 0 | 0 |
| TM027 | w/o female | RS: 2, FS: 1 | RS: 12, FS: 0 | RS: 1, FS: 1 | 0 | 1 | 2 | 0 | 0 | 2 | 0 |
| TM028 | w/ female | RS: 8, FS: 0 | RS: 4, FS: 2 | RS: 11, FS: 1 | 3 | 0 | 0 | 2 | 0 | 0 | 0 |
| TM029 | w/o female | RS: 14, FS: 0 | RS: 7, FS: 0 | RS: 7, FS: 0 | 4 | 0 | 0 | 0 | 1 | 0 | 0 |
| TM033 | w/ female | RS: 5, FS: 1 | RS: 11, FS: 1 | RS: 4, FS: 0 | 2 | 1 | 1 | 0 | 0 | 1 | 0 |
| TM035 | w/o female | RS: 0, FS: 0 | RS: 3, FS: 0 | RS: 5, FS: 3 | 0 | 0 | 1 | 0 | 0 | 0 | 2 |
| TM037 | w/ female | RS: 5, FS: 1 | RS: 20, FS: 0 | RS: 13, FS: 3 | 4 | 0 | 5 | 0 | 0 | 0 | 0 |
| TM038 | w/o female | RS: 3, FS: 0 | RS: 3, FS: 1 | RS: 1, FS: 1 | 0 | 0 | 0 | 0 | 0 | 1 | 0 |
| TM039 | w/o female | RS: 8, FS: 1 | RS: 6, FS: 2 | RS: 10, FS: 2 | 5 | 0 | 0 | 0 | 0 | 0 | 1 |

